# High-throughput optimisation of protein secretion in yeast via an engineered biosensor

**DOI:** 10.1101/2024.05.15.594099

**Authors:** Alexandra Williams, Runpeng Luo, Oliver B. Smith, Lydia Murphy, Benjamin Schwessinger, Joseph Brock

**Author notes:** Correspondence (J.B.).

## Abstract

Secretion of high value proteins and enzymes is fundamental to the synthetic biology economy; allowing continuous fermentation during production and protein purification without cell lysis. Most eukaryotic protein secretion is encoded by an N-terminal signal peptide; however, the strong impact of signal peptide sequence variation on the secretion efficiency of a given protein is not well defined. Despite high natural signal peptide sequence diversity, most recombinant protein secretion systems employ only a few well characterised signal peptides. Additionally, the selection of promoters and terminators can significantly affect secretion efficiency, yet screening numerous genetic constructs for optimal sequences remains inefficient. Here, we have adapted a yeast G-protein coupled receptor biosensor, to measure the concentration of a peptide tag that is co-secreted with any protein of interest. Protein secretion efficiency can thus be quantified via the induction of a fluorescent reporter that is upregulated downstream of receptor activation. This enables high-throughput screening of over 6000 combinations of promoters, signal peptides and terminators, assembled using one-pot Combinatorial Golden Gate cloning. We demonstrate this biosensor can quickly identify best combinations for secretion and quantify secretion levels.

**Graphical Abstract:** 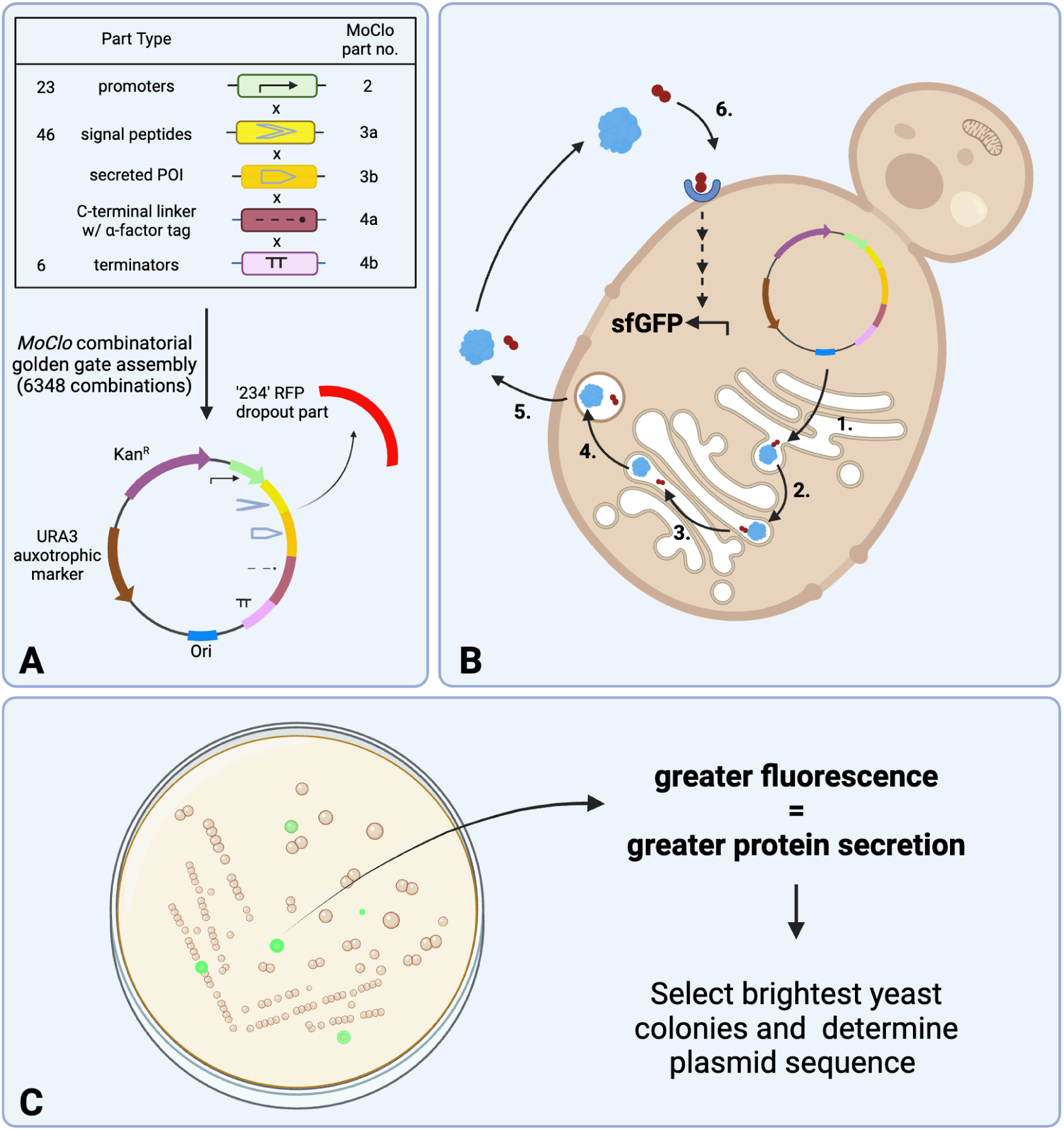

## Introduction

Bakers yeast (*Saccharomyces cerevisiae)* has been used for thousands of years by human societies to increase the value and digestibility of grain. ^1^ More recently, yeast has become an industrially important synthetic biology chassis for the production of high value recombinant proteins. *S. cerevisiae* and other yeast species are ideal expression hosts for recombinant Eukaryotic proteins, as they can perform post-translational modifications necessary for protein function. ^2^ Valuable recombinant proteins currently produced in yeast include peptide-based biopharmaceuticals, industrial biocatalysts and proteins for lab-grown food alternatives. ^2–8^

When producing recombinant proteins for industrial use, it is advantageous if the protein of interest (POI) can be secreted from the expression host. ^9^ Secretion of the POI allows for continuous precision fermentation during production and avoids the need for cell lysis, which is uneconomical at scale. Optimising recombinant protein sequences for secretion can thus result in more efficient and profitable manufacturing of valuable target proteins/enzymes. ^2, 7^

Signal Peptides (SPs) play an important role in delivering ribosome-nascent proteins to the endoplasmic reticulum (ER) membrane, and the subsequent translocation of the proteins into the ER lumen; the first step in the eukaryotic protein secretion (SEC) pathway. ^10^ SPs typically consist of a series of hydrophobic amino acids, flanked by a positively charged N-terminal domain and a polar, uncharged C-terminal domain which includes a signal peptidase cleavage site. ^10, 11^ When the SP is cleaved, this releases the secreted protein into the ER lumen for trafficking into the SEC pathway via the Golgi apparatus. ^12^ Secretory proteins can enter the ER during (co-translational secretion) or after translation (post-translational secretion). While co-translational secretion is largely a passive process relying on Brownian motion for translocation, post-translational secretion requires cytosolic and luminal chaperones for translocation and folding, resulting in significant nucleoside triphosphate turnover. ^10^ The size and hydrophobicity of the signal peptide and C-terminal amino acid sequence, influence which pathway is taken, with some proteins able to access both pathways. ^10, 12^ SPs that direct for ‘post-translational translocation’ commonly contain a ‘pre-pro’ structure. The ‘pre’ region roughly corresponds to the conventional SP structure described previously, and the ‘pro’ region contains internal protease cleavage sites for subsequent processing during the SEC pathway. ^10, 11^ ‘Pro’ region sequences can vary greatly in sequence, but are essential for translocation into the ER, interacting with molecular chaperones which can aid in proper protein folding (**Figure S1**). ^13, 14^

At present, most recombinant proteins are engineered to be secreted from yeast using just a few common SPs. ^10, 15^ However, this strategy does not always lead to successful POI secretion. The efficiency of signal peptidase cleavage is influenced by both the upstream residues (indicated by negative indices) of the SP and the downstream residues (indicated by positive indices) of the POI. ^16–18^ Thus, a SP that is optimal for one protein might not be as effective for another. Evolution has selected each SP for its ability to efficiently secrete its corresponding endogenous protein within its source organism. ^19, 20^ For heterologous expression, POI secretion can be optimised by testing different combinations of promoters and terminators concurrently with SPs,^4, 9^ that collectively form a library of ‘protein expression cassettes’. This approach can yield significant and unpredictable increases in expression yields. ^4, 21^ For example, it has been demonstrated that a strong promoter will sometimes result in less efficient secretion than a weaker promoter for some SP+POI combinations,^22^ since overloading of the secretory machinery results in an unfolded protein response within the ER. ^23, 24^ It is therefore essential that SP+POI sequence pairing is optimal to achieve the most efficient protein secretion. ^2, 4, 7–9, 11, 22, 25^ While some progress has been made in bioinformatically predicting the efficiency of particular signal peptides for recombinant target proteins,^26–28^ these tools are in the early stages of development and lack means of high-throughput validation screens. The most reliable optimisation method is often via an empirical trial and error process of testing different SP sequences one by one. The process of individually testing several SPs^14^ is often time consuming, costly and inefficient. Notably there are several existing protein secretion screening systems relying on chromogenic or fluorogenic substrate assays, split-GFP or larger enzymatic tags for cell survival selection. While making significant progress on this problem, these methods still have drawbacks such as being substrate-dependent, labour intensive, or require large tags which may limit POI secretion efficiency. ^4, 8, 9, 26, 29–31^

This highlights the need for a rapid and direct way to screen the secretion efficiency of any POI in a near native state, when assembled into a high-diversity SP and expression cassette library. Screening such diverse expression cassette libraries requires a high-throughput method to select the best cassette combinations for POI secretion. To achieve this, we built upon the work of Shaw *et al.,* who previously engineered G-protein coupled receptor (GPCR)-based living biosensors in *S. cerevisiae* by refactoring a chassis strain with non-essential components of the signalling cascade removed (yWS677). ^32^ This allowed for genetic refactoring to give linear, orthogonal signalling from receptor activation to reporter gene expression. Critical for orthogonality was the use of a synthetic transcription factor and modular promoter, allowing for creation of highly specific living biosensors with tuneable dose response capabilities. We aimed to adapt this system for measuring protein secretion efficiency via the native Ste2 receptor, for which the endogenous agonist is the hexapeptide alpha-mating factor (ɑMF). ^33^ We have developed a whole-cell living biosensor for protein secretion, using a minimal cleavable peptide receptor agonist tag that is automatically cleaved and co-secreted with the POI (**Figure 1**). We quantified the diversity of our library via long read Nanopore sequencing (**Figure 2**) and benchmarked this system using recombinant Human Serum Albumin (HSA). Secretion of HSA from *S. cerevisiae* is well documented in the literature. ^34, 35^ We rapidly identified new genetic combinations with comparative secretion efficiency to the native signal HSA signal peptide. We found our secretion biosensor shows an excellent linearity of dose-response (**Figure 3 and 4**).

**Figure 1.**
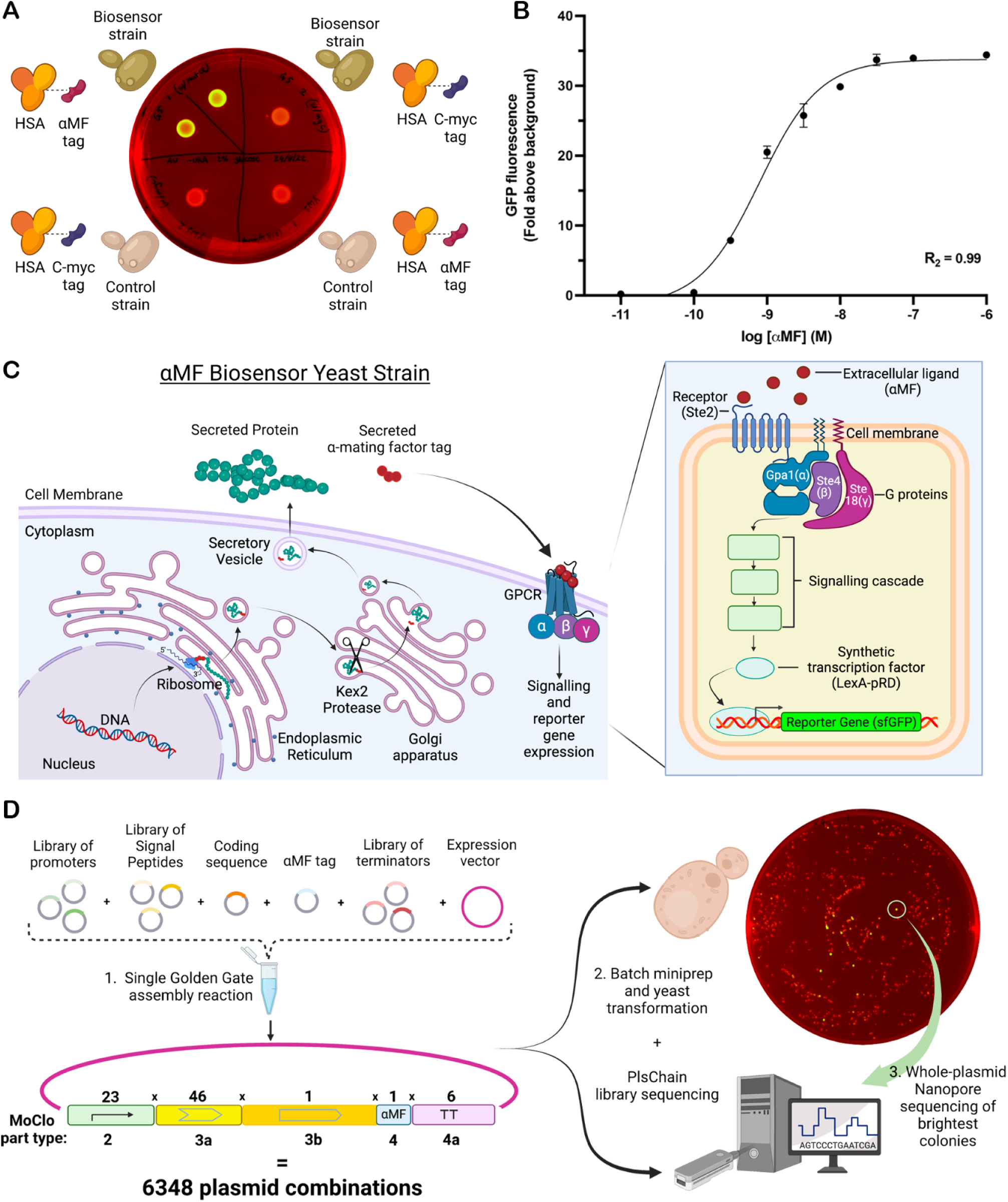
Biosensor assembly and workflow. A) An agar plate, inoculated with 20 μL spot cultures of yeast cultures at equal cell densities was incubated at 30 ℃ for 48 hours and imaged separately for green fluorescent emissions and under white light for background. These images were subsequently merged to effectively differentiate variations in sfGFP expression, with false colouring applied in green and red for each image type, respectively. Specificity of the biosensor was validated through a comparative analysis. An αMF-tagged (WHWLQLKPGQPMY) Human Serum Albumin (HSA) construct, under its native signal peptide and a strong promoter, was transformed into the biosensor strain (displayed in the top left quadrants). This was contrasted with the transformation of the same construct into a control strain yWS677 (illustrated in the bottom right quadrant). Additionally, to confirm the absence of cross-sensitivity, the biosensor strain was transformed with an HSA construct tagged with a c-myc peptide (AAAEQKLISEEDL) instead of αMF (shown in the top right quadrant). It was observed that this construct did also not induce fluorescence in the control strain (noted in the bottom left quadrant). B) A Biosensor operational range of ∼3 orders of magnitude (∼0.1 - 100 nM) was demonstrated by creating a dose response curve. Increasing concentrations of synthetic ɑMF peptide was titrated into aliquots of log-phase liquid culture and the average induced fluorescence measured by flow cytometry. Each data point represents Mean ± Standard Error of the Mean (SEM), n = 3 technical replicates, each containing ≥20000 biological replicates (single cells). Curve span = 35.21. Curve parameters: Bottom (a) = −1.448, Top (d) = 33.77, EC50 (c) = 7.869e-10, HillSlope (b) = 1.052. Values represented are the fold increased fluorescence relative to the control strain. Figure made using Prism 9 (GraphPad). C) An overview of the biosensor design showing translocation of a secreted POI with an ɑMF C-terminal tag that is cleaved by Kex2 in the late golgi, before being co-secreted (left) and its subsequent activation of the Ste2 and downstream orthogonal signalling to activate the synthetic transcription factor leading to reporter gene (*sfGFP*) expression (right). D) Overview of combinatorial Golden Gate assembly theoretically creating >6000 unique combinations that can be verified using PlsChain analysis of Nanopore library sequencing after performing a batch miniprep of transformed *E. coli* (Figure 2D). The same DNA library can then be transformed into yeast, and the most fluorescent colonies sequenced to determine the optimal SP expression cassette for the POI using whole plasmid Nanopore sequencing. Agar plate of transformed library in yeast imaged as in (A).

**Figure 2:**
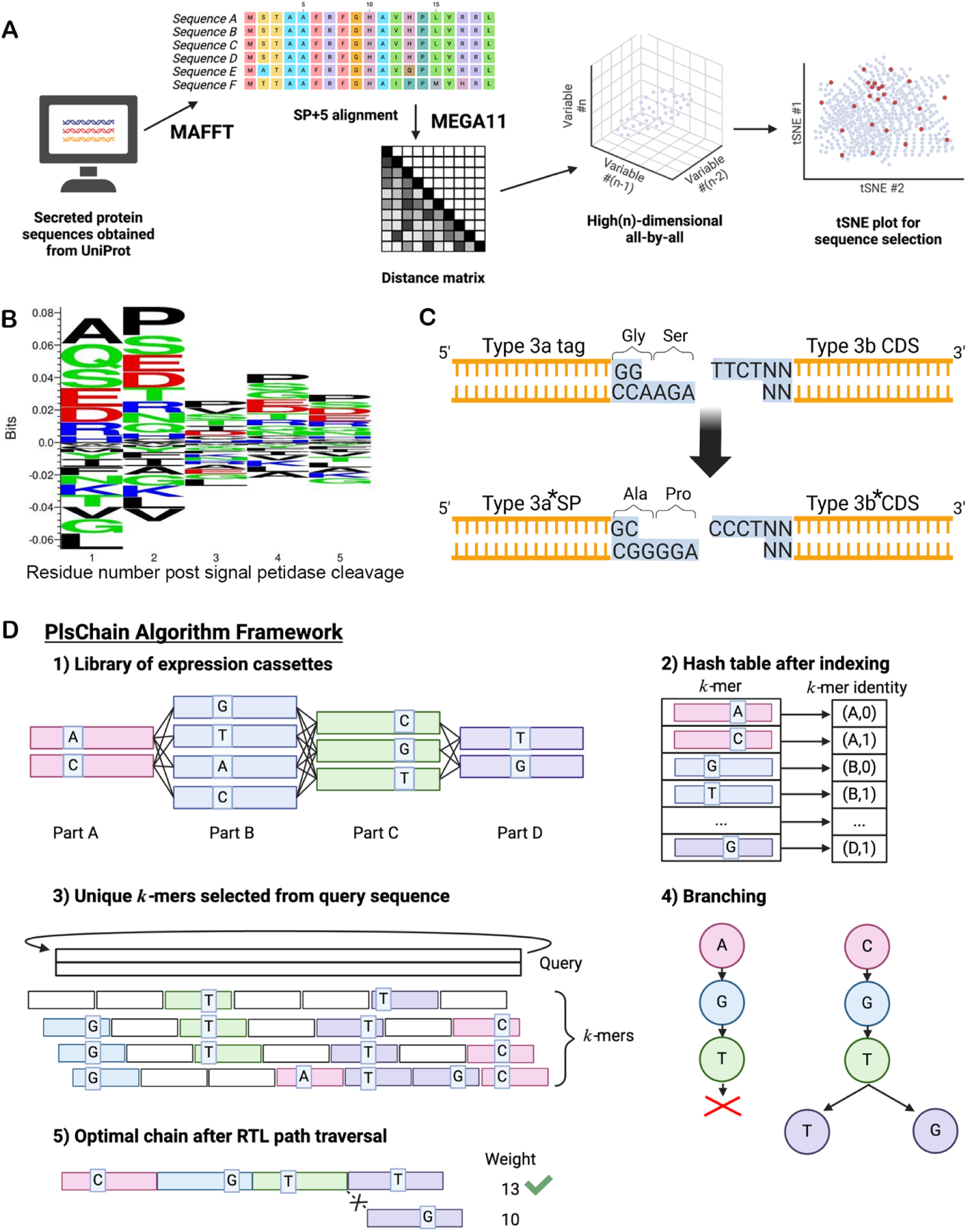
**MoClo Signal peptide library design** A) Workflow for analysis of signal peptide datasets. Initially, all SwissProt proteins in UniProt from *H. sapiens* and *S. cerevisiae* that were annotated as containing an N-terminal signal peptide were downloaded and processed to produce independent datasets of SP+5 sequences (signal peptide + the following five residues) for each species. MAFFT was used to produce a multiple sequence alignment of each dataset, followed by MEGA11 to produce an all-by-all distance matrix based on each multiple sequence alignment. The dimensionality of this high (n)-dimensional all-by-all representation was reduced using t-SNE to produce a Cartesian plot from which a maximally diverse set of signal peptides (red) could be selected based on maximising the Euclidean distance between selected signal peptides compared to the background, non-selected signal peptides (blue). B) SequenceLogo^44^ analysis of all Eukaryotic signal peptides +5 residues downstream of the signal peptidase cleavage site predicted by Signal P, ^41^ showing a preference for Alanine and Proline at the +1 and +2 positions, respectively. A similar enrichment was observed when analysing the *S. cerevisiae* and *H. sapiens* datasets separately (Figure S2). C) Design of the 3a-3b junction of the MoClo assembly standard (above) within our signal peptide library (below) to introduce an Alanine and Proline at the +1 and +2 positions, respectively. D) The PlsChain algorithm framework. 1) An example for a library of expression cassettes with 4 components A,B,C,D labelled in different colours, variations within each component is identified by rectangular boxes. 2) Index is constructed based on k-mer counting from the group of components. 3) unique k-mers are identified from the query sequence and labelled in specific colours. k-mers represented by hollow rectangles are non-unique. 4) Branchings are constructed and RTL (Register Transfer Level) path traversal is performed. 5) An optimal chain with weight 13 is selected to denote the class of Query. Figure made using BioRender.com.

**Figure 3.**
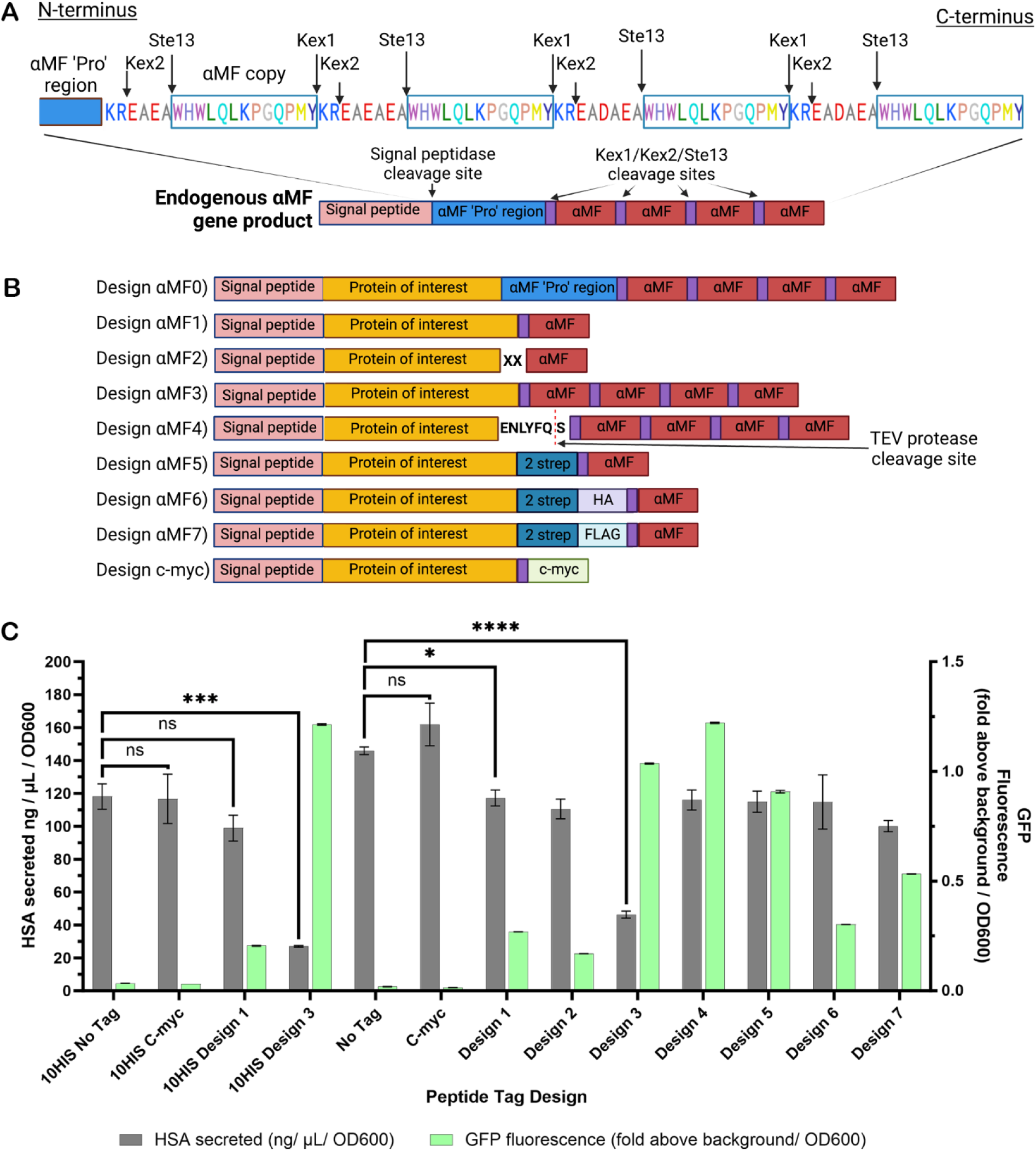
**ɑMF biosensor yeast strain responds to synthetic and self-secreted S. cerevisiae ɑMF peptide:** A) Native S. cerevisiae ɑMF pre-peptide sequence with Kex1, Kex2 and Ste13 proteolytic cleavage (above) and Schematic of C-terminal ɑMF peptide tag designs (below). Kex2 cleaves after the KR sequences to separate the four copies of ɑMF. Ste13 cleaves the EA sequences from the N-terminus of the aMF copies. Kex1 cleaves the KR sequence from the C-terminus of the first three aMF sequence copies. Figure made using Biorender.com. B) C-terminal peptide tag designs for use with ɑMF biosensor. Design ɑMF0) Full native *S. cerevisiae* ɑMF sequence minus the ‘Pre’ region but including the ‘Pro’ region and four ɑMF peptide repeats. Design ɑMF1) One copy of the ɑMF peptide with the N-terminal native Kex2 and Ste13 protease cleavage sites present to allow for cleavage from the POI during the secretion pathway. Design ɑMF2) One copy of the ɑMF peptide directly conjugated to the C-terminus of the POI, missing the Kex2 and Ste13 protease cleavage sites (no KREAEA sequence). Design ɑMF3) the four native copies of the ɑMF peptide, with all protease cleavage sites intact, but no ‘Pro’ region. Design ɑMF4) One copy of the ɑMF peptide with a N-terminal TEV protease cleavage site. Design ɑMF5) Twin-strep purification tag directly conjugated to the C-terminus of the POI, followed by one copy of the ɑMF peptide with native protease cleavage sites intact. Designs ɑMF6-7)Analogous to Design ɑMF5 but with a HA and FLAG tag after the protease cleavage sites, respectively. Design c-myc) analogous to Design ɑMF1, but with a c-myc-tag in place of the ɑMF peptide as a negative control. Figure made using Biorender.com. C) Comparison of C-terminal ɑMF tag design for biosensor activation (mean single cell GFP fluorescence, fold above background/OD600) versus secreted HSA productivity (ng/ml/OD600) of 72 hr *S. cerevisiae* (biosensor strain) HSA-tag expression cultures. 10HIS refers to the HSA expression sequence with an N-terminal 10xHIS tag. Brackets indicate multiple t test statistical comparisons: ns = not significant (p-value >0.05), * = <0.05, ** = <0.01, *** = 0.005, **** = <0.0001. Graph made and analysis conducted using Prism GraphPad.

**Figure 4.**
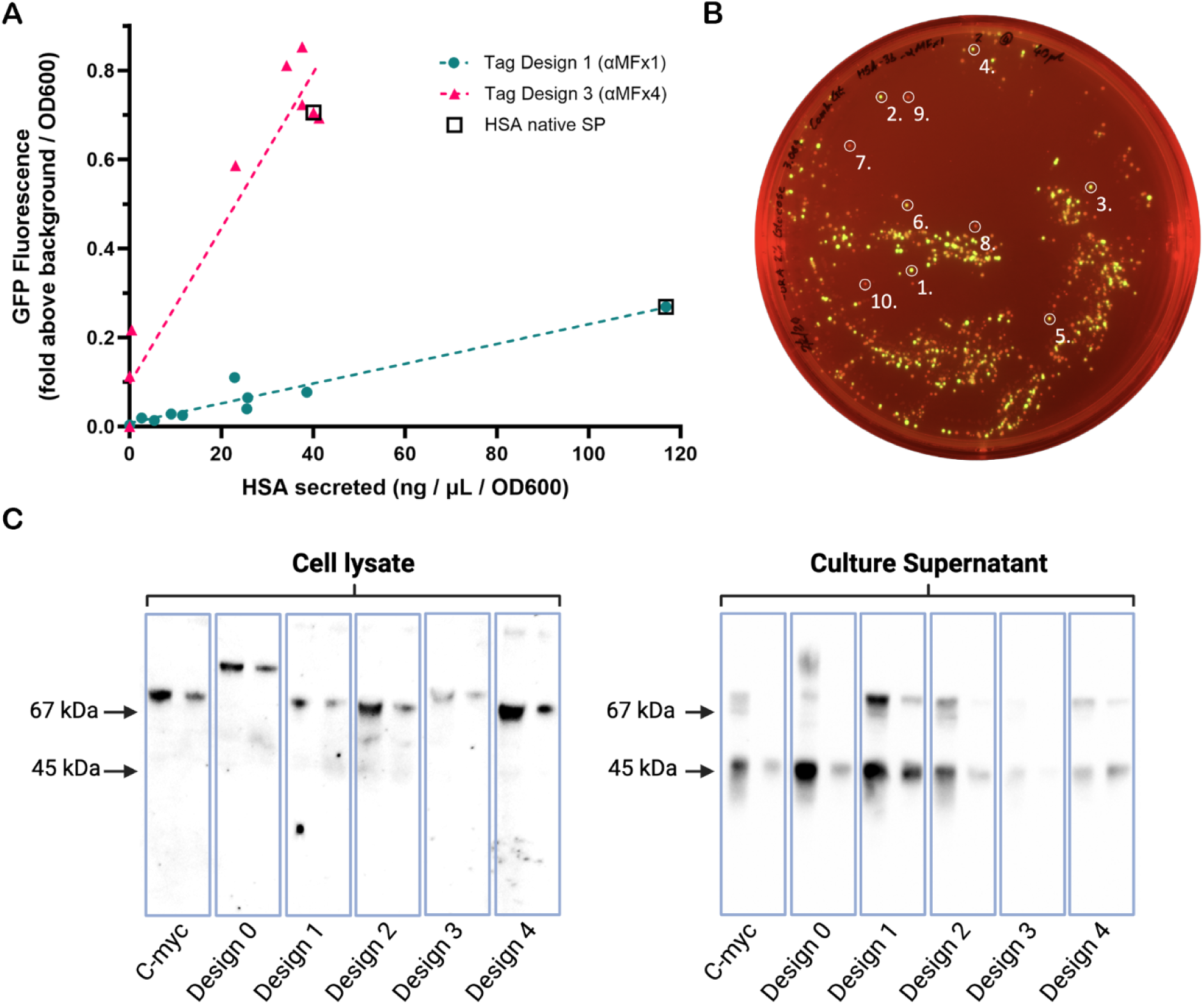
**Validation of the secretion biosensor with combinatorial assemblies of HSA and different ɑMF tag designs** A) Comparative analysis of biosensor GFP response (RFU fold above background/OD600) vs HSA secreted (ng/mL/OD600). Each point represents a sample with a different HSA expression cassette combination. Blue circles show samples assembled with tag Design ɑMF1, and pink triangles show samples assembled with tag Design ɑMF3. Both data sets follow linear trendlines, with y=0.017*x+0.096, R^2^ = 0.92 and y = 0.0022*x+0.0075, R^2^ = 0.93 for Design ɑMF3 and Design ɑMF1, respectively. Square boxes highlight samples assembled with the native HSA signal peptide sequence. B) Example plate image of ɑMF biosensor yeast strain transformed with Combinatorial library of HSA assembled with tag Design ɑMF1 expression cassettes. C) Western blot results comparing cell lysate and culture supernatant from secretion biosensor strain 72 hr HSA expression cultures, demonstrating cleavage of c-terminal ɑMF. Antibodies used for Western blot: mouse anti-HSA monoclonal primary antibody (Invitrogen) and Horseradish peroxidase (HRP) conjugated goat-anti-mouse secondary antibody (BioRad). Expected size of HSA without c-terminal tag is ∼67 kDa. A well characterised HSA degradation product is visible at ∼45 kDa.

## Results and Discussion

To efficiently identify the best SP for a given POI, we first developed a set of 46 maximally diverse SPs by analysing alignments of whole genome-wide SP sequences (**Figure 2**). We analysed SP sequences predicted to target the co-translational pathway only, since it has been shown this can lead to increases in secretion of heterologous targets and is hypothesised to impart a lower metabolic stress on the cell. ^10, 12, 14^ These sequences were then designed for high-throughput generation of protein expression cassette plasmids, and combined with a well characterised library of promoters and terminators of the MoClo Yeast Toolkit (YTK)^36^ using Combinatorial Golden Gate cloning. ^4, 21, 37, 38^ Combinatorial Golden Gate assembly utilises a library of modular cloning parts, with the same flanking overhang sequences when digested with a type IIS restriction enzyme (RE), such that only one part from the library is assembled at random into a given position in each plasmid. ^21^ Combinatorial optimisation is a common strategy in synthetic biology^39^ and others have also applied this approach to effectively optimise protein secretion. ^4, 8, 21^ Here we extend on this approach by showing that Combinatorial Golden Gate assembly using our library of diverse signal peptides together with the MoClo YTK parts library^36^ can generate over 6000 unique protein expression cassette combinations in a single reaction. Importantly, we developed a Nanopore long-read sequencing approach that validates the completeness and evenness of our screening library. This is a significant progress because previously these factors were only assumed. To the best of our knowledge, this is the first time combinatorial Golden Gate cloning has been demonstrated and verified at this scale, which we have quantified using a novel algorithm, PlsChain, to efficiently identify unique sequence combinations (**Figure 2D**).

The native *S. cerevisiae* ɑMF gene sequence consists of a ‘pre-pro’ secretion signal and four tandem ɑMF copies separated by spacers that are cleaved by multiple late-ER and golgi localised proteases (Kex1, Kex2 and Ste13) during trafficking through the SEC pathway (**Figure 3A**). ^33, 40^ By including a C-terminal Kex2 protease sequence followed by an ɑMF peptide tag on the POI coding sequence during combinatorial assembly, we hypothesised that it would be cleaved and co-secreted alongside the POI (**Figure 1C**). Thus, the secreted ɑMF peptide could activate the Ste2 receptor and stimulate expression of a sfGFP reporter gene, effectively coupling POI secretion efficiency to yeast colony fluorescence. The brightest colonies could then be selected and sequenced to determine the best protein expression cassette combinations for efficient POI secretion (**Figure 1D**). We subsequently designed, tested and optimised this biosensor using various tag designs for protein secretion using the yWS677 chassis strain from Shaw *et al.* (**Figure 3**). ^32^

### Designing a maximally diverse signal peptide library

To generate a diverse set of SPs to augment the MoClo YTK,^36^ we undertook a bioinformatic analysis of eukaryotic secreted proteins. We first conducted a search using UniProt for experimentally characterised secreted proteins with SPs from *Saccharomyces* and *H. sapiens,* which were of particular interest due to their relevance to the expression host and the production of biopharmaceuticals with correct folding and function, respectively. We then processed these sets of protein sequences using SignalP 5.0 to produce sequence sets of distinct “SP+5” sequence sets, including five residues downstream of the signal peptidase cleavage site predicted by the software. ^41^ By including an additional five residues beyond the cleavage site, we aimed to conduct a weighted sequence analysis on both the SP sequences and the cleavage site itself, catering to signal peptidase processing. ^16–18^ MAFFT^42^ was used to independently align each sequence set, followed by MEGA11^43^ to generate a dataset-specific all-by-all distance matrix with expectation maximisation. Given this method provides high-dimensional embedding of the diversity in each set of SPs, we used single value decomposition (SVD) followed by t-distributed stochastic neighbour embedding (t-SNE) to visualise the diversity based on relative positioning in a three-dimensional plot. After multiple iterations for consistency, the resulting clusters in the t-SNE plots reinforced the notion of iterative evolutionary divergence among signal peptides. This provided a means to identify divergent SPs based on Euclidean distance to other SPs in the plot. We selected 24 SP sequences from *Saccharomyces* and 22 SP sequences from *H. sapiens* that were maximally distant to create a SP library **(Figure 2A)**. Finally, we analysed the aligned SP+5 sequences of all Eukaryota secreted proteins (**Figure 2B)**, in addition to the *Saccharomyces* and *H. sapiens* in isolation (**Figure S2**), using SequenceLogo,^44^ to find the optimal sequence introduced by the MoClo cloning junction. We found a qualitative enrichment of an Alanine and Proline residue at positions, +1 and +2 downstream of the predicted cleavage site, respectively. We subsequently redesigned the type 3a (SP) to 3b (POI) junction to encode an Ala-Pro rather than the Gly-Ser defined by the MoClo assembly standard (**Figure 2C**). ^36^

Following synthesis of this unique SP library as MoClo compatible type 3a* parts, these were combined with Human Serum Albumin (HSA) synthesised as a type 3b* part (* denotes that the MoClo part contains the Ala-Pro 3a-3b junction) as a test POI, since it provided a useful benchmark for protein secretion efficiency in yeast. ^34, 45, 46^ These parts were combined with one of several synthesised ɑMF type 4a C-terminal tags (**Figure 3B**), in addition to the library of well characterised promoters (type 2) and terminators (type 4b) provided in the MoClo YTK (**Figure 1D**). ^36^

### Optimising ‘Combinatorial’ Golden Gate assembly of protein expression cassettes

Golden Gate assembly has been used to assemble up to 52 component parts in tandem by optimising overhang ligation fidelity, via a data-optimised assembly design (DAD) algorithm. ^47^ Combinatorial Golden Gate assembly, however, has been less rigorously characterised. For example, Püllmann and Weissenborn ^4^ have demonstrated combinatorial Golden Gate assembly to optimise protein secretion in yeast, using 187 possible combinations of parts. However, these authors and others have not quantified the success and evenness of such combinatorial libraries, despite the importance for quality control purposes.

Theoretically, the maximum number of possible combinations that can be generated using Combinatorial Golden Gate assembly is limited only by the number of genetic parts used in assembly and the transformation efficiency of the recipient organism. We hypothesised that this technique could be used to assemble thousands of protein expression cassette combinations in a single reaction. Using 23 promoters and 6 type 4b terminators from the MoClo YTK,^36^ along with the 46 diverse signal peptides, a chosen POI and ɑMF c-terminal tag, we aimed to assemble 6348 protein expression cassette combinations, in a single Combinatorial Golden Gate assembly reaction **(Figure 1D)**. Following assembly, the Combinatorial Golden Gate ligation mixture was transformed into *E. coli* to select for intact plasmid assemblies and to amplify the copy number of each expression cassette. Combinatorial POI expression cassette libraries were then extracted from *E. coli* via batch plasmid miniprep. The resulting POI expression cassette plasmid libraries were assessed via nanopore long-read sequencing.

To efficiently assess whether assembly of the complete expected library of expression cassettes (6348 combinations) was being achieved, we developed PlsChain. This new classifier algorithm employs the strategies of unique *k*-mer identification and branching^48^, to index the library of expression cassettes and classify the reads into one of the 6348 possible combinations (**Figure 2D**). We benchmarked PlsChain against the minimap2^49^ alignment method on simulated datasets. This benchmarking demonstrates that PlsChain achieves the best precision and recall rate while maintaining low computational costs (see, **Figure S4-S5**, **Table S6** and methods for detail).

We analysed three different library assemblies of HSA expression constructs. Sequencing results classified by PlsChain revealed that ∼95 % (mean of 2 biological replicates, each with 3 technical repeats ∼95% = 6028 combinations) of the total expected combinations were represented in our expression cassette libraries (see **Supplementary File 2** for high resolution heat map visualisation and tabulation of reads versus sequence combinations). Based on the results we found that some expression cassette combinations were only represented once in the fully classified library, while others were represented >500 times. Heat map visualisation also revealed preferences for certain promoter and signal peptide combinations. This bias toward particular part sequences arranging together could be due to variations in part length, unevenness in the batch miniprep, or other experimental error. We found that ensuring precise equimolarity of all parts was essential for successful assemblies. In future experiments, error could potentially be reduced by using a more accurate method for determining DNA concentrations of parts, such as a Qubit fluorometer, and automated liquid handling robotics for setting up assembly reactions. Our results however, show that the randomised combinatorial assembly was sufficient for testing over 6000 of the possible 6348 expression cassette combinations in high-throughput using our secretion biosensor.

### Building and testing a biosensor for protein secretion

To create the *S. cerevisiae* ɑMF biosensor we used components from the MoClo YTK and the GPCR toolkits. ^32, 36^ Using the Level 0 components from these kits we assembled: Level 1 gene assemblies for the ɑMF biosensor receptor (*Ste2*), G-protein alpha-subunit (*Gpa1*), synthetic transcription factor (*LexA-PRD*) and reporter gene (*sfGFP*) with modular *LexO* promoter via Golden Gate assembly; along with a markerless integration vector with ‘234’ *GFP* dropout site and *LEU2* site homology arms, designed to be compatible with the *LEU2* landing site in the yWS677 strain to mimic the ‘design 4’ biosensor from Shaw *et al.* ^32^ The Level 2 construct, *LEU2* integration cassette containing the 4 biosensor genes, was assembled via Golden Gate assembly **(Figure S3)**. Plasmid sequences were verified at each level via nanopore whole plasmid sequencing.

Following sequence confirmation of the Level 2 integration cassette containing the biosensor genes, the genes were inserted into the host yeast genome via CRISPR/Cas9 mediated gene editing. Single colonies were selected and screened for successful gene insertion via gDNA extraction and a series of PCR reactions. The results of the diagnostic PCR reactions were consistent with the expected band profile for successful integration of the biosensor gene cassette. Thus we were able to continue on to testing the activity of the biosensor strain in response to the ligand *S. cerevisiae* ɑMF **(Figure S3)**.

Initially the potential ɑMF biosensor strain was screened for biosensor activation by applying water (-ve control) or 100 nM synthetic *S. cerevisiae* ɑMF peptide. Based on the post hoc reverse pairwise comparisons performed in R, there was no significant difference in fluorescence between the ‘Water’ and ‘ɑMF’ treatments of the control yeast strain (yWS677) and there was a highly significant difference between the ‘Water’ and ‘ɑMF’ treatments of the aMF biosensor yeast strain (**Supplementary file 3**). Based on the Two-way ANOVA performed in R, there was a highly significant (p≤0.001) difference between the variances in RFU by treatment and yeast strain; indicating that the aMF biosensor responds to treatment with synthetic *S. cerevisiae* ɑMF peptide by producing green fluorescence (**Figure S3D**). Following initial confirmation of biosensor activation via addition of synthetic *S. cerevisiae* ɑMF, dilutions of ɑMF were prepared and applied to samples of the biosensor strain. To generate an ɑMF biosensor dose-response curve, single cell GFP fluorescence was measured via flow-cytometry. The lowest fluorescence signal readings observed were ∼0.2 fold over background at [10^-11^] M ɑMF added to the sample cultures. The highest fluorescence signal readings observed were ∼34 fold over background at [10^-5^] M ɑMF added to the sample cultures (**Figure 1B**).

To determine if the engineered biosensor strain could self-activate the biosensor pathway through the secretion of a recombinant protein tagged with a C-terminal ɑMF peptide tag, we chose recombinant Human Serum Albumin. HSA has been routinely secreted from *S. cerevisiae* in published literature, and studies have shown that changing the associated SP sequence can lead to differences in HSA secretion efficiency. ^34, 45, 46^ This made HSA an ideal test case for our system. We designed a codon optimised HSA coding sequence as a compatible type 3b* part (**Figure 2C**) with and without a 10xHis N-terminal tag. For the initial HSA expression constructs, we used the strong constitutive promoter pCCW12 (Type 2), the native HSA SP sequence (Type 3a*), the 10xHis-HSA coding sequence (Type 3b*) and the tDH1 terminator (Type 4b). With this common set of parts, we systematically evaluated a series of C-terminal tag configurations (Type 4a), progressively simplifying their structure. These designs, derived from the endogenous *MF(ALPHA)1* gene of *S. cerevisiae* (**Figure 3A**), aimed to delineate the minimal peptide tag configuration that would minimise any adverse effects on protein folding and secretion, while still triggering a quantifiable response in the biosensor (**Figure 3B**, full sequences in **Supplementary file 4**). All tags were then tested in parallel to assess their efficacy. We developed eight distinct ɑMF tag configurations, beginning with the full coding sequence post-’Pre’ region (Design ɑMF0) and progressively simplifying to a singular ɑMF motif both with native Kex2/Ste13 protease sites intact and a non-cleavable variety for comparison with this recognition sequence replaced with a glycine-serine linker sequence. A C-terminal c-myc tag as well as a construct with no c-terminal tag were used as negative controls; hypothesised to lack any activating influence on the αMF biosensor pathway. Figure 3C compares the mean single cell GFP fluorescence (Fold above background, quantified via flow cytometry), with secreted HSA (ng/mL, quantified via HSA ELISA), both normalised to culture optical density at 600 nm (OD600), for biosensor yeast strain cultures expressing HSA with various C-terminal ɑMF tag designs (**Figure 3C)**.

### Combinatorial library production with a ɑMFx4 tag

Initially, 4 ɑMF tag designs (ɑMF0 - ɑMF4) were confirmed to activate the ɑMF biosensor strain via co-secretion of HSA with a C-terminal ɑMF tag via a spot plate assay (design ɑMF3 shown in **Figure 1A**). We then performed Western blot analysis that showed successful secretion of the HSA constructs containing tag designs ɑMF0-ɑMF4 at a size of ∼67 kDa. Although the molecular weight (MW) change, corresponding to cleavage of the C-terminal tag, was too small to observe for designs ɑMF1-2 (ɑMFx1 tags); successful cleavage of design ɑMF0 and ɑMF3 containing 4 copies of the ɑMF peptide was clearly evident by comparing the MW difference of these samples from cell lysate (uncleaved, MW ∼79 kDa) with those of the cell supernatant (cleaved, ∼67 kDa). While Design ɑMF0 containing the native ‘pro’ region followed by 4 repeats of the *MF(ALPHA)1* gene showed robust biosensor activation, western blotting showed heterogeneity in the MW of the secreted protein. Based on these results and qualitative comparisons of flow cytometry fluorescence, Design ɑMF3 (ɑMFx4 tag) appeared to give a robust biosensor response that correlated with efficient protein secretion. Initial Combinatorial Golden Gate assembly experiments with all promoters, SPs and terminators, were conducted using this tag. Colonies displaying dull, medium and bright colony fluorescence were selected and used to inoculate liquid cultures whose fluorescence and HSA secretion could be accurately compared using flow cytometry and a HSA ELISA (enzyme-linked immunosorbent assay) of the culture supernatant, respectively. Colonies from a combinatorial library transformation corresponding to Design (ɑMF3) were subsequently chosen to quantitatively determine HSA secretion. Identical constructs containing the native HSA SP and Designs ɑMF1-4, c-myc and no-tag negative controls were also included in the ELISA to compare the effect of tag design on protein secretion efficiency. All cultures were assayed in triplicate and corrected for variations in culture density (**Figure 4A** shown in pink and **Supplementary File 4**). Results showed that we were able to rapidly identify novel constructs that achieved comparable secretion efficiency to the native SP positive control. It was clear however, that the Design ɑMF3 tag was having a significant (*p <* 0.05) impact on HSA secretion compared to the c-myc and ‘No tag’ negative controls (**Figure 3C**). In contrast however, Design ɑMF1 had a small negative effect on protein secretion while still achieving a measurable biosensor response above baseline.

### Combinatorial library production with a ɑMFx1 tag

Having completed one iteration of the Design-Test-Build-Learn cycle, we identified the simplest tag design (ɑMF1) as having the smallest effect on protein secretion efficiency while still activating the biosensor. We subsequently generated designs ɑMF5-7 from design ɑMF1 to also include affinity and epitope tags for purification and non-POI dependent immunohistochemistry, respectively (**Figure 3C**). A second round of Combinatorial Golden Gate assembly experiments was then performed using the Design ɑMF1 tag (ɑMFx1 tag) (**Figure 4A,** shown in green). For this analysis, we also chose to perform assemblies with a HSA genetic part (type 3b*) lacking a N-terminal His-tag, to more accurately reflect the ability of our signal peptide library diversity to enhance secretion by complementing the native N-terminal sequence of the POI. ^16–18^ As before, colonies displaying varying levels of colony fluorescence were selected for comparison using flow cytometry and a HSA ELISA of the culture supernatant, respectively. During this second round of testing, a focus was placed on colonies with intermediate fluorescence, since these data points were underrepresented in our analysis of the ɑMF3 tag assembly (**Figure 4A** shown in pink and **Supplementary File 5**). Designs ɑMF5-7 all contained a twin-strep tag, linked to the C-terminus of the POI, followed by a single ɑMF copy. Design ɑMF6 and ɑMF7 also contained an interleaving HA (Human influenza hemagglutinin) and a FLAG tag, respectively (**Figure 3b**). These constructs were also compared using flow cytometry and HSA ELISA analysis as above. Each of these HSA ɑMF tag test constructs used the same promoter (pCCW12), the native HSA SP sequence, the non-His-tagged HSA coding sequence and the tDH1 terminator, so that the effect of different tag designs on secretion efficiency could be objectively quantified (**Figure 3C**). In this analysis, we detected a small but statistically significant (*p <* 0.05) negative effect of the ɑMF1 tag on protein expression efficiency relative to the no-tag control (**Figure 3C**).

### Sensitivity and dynamic range of the secretion biosensor is altered using different ɑMF tag designs

In each of the experiments (using Design ɑMF3 and Design ɑMF1), we included a construct with the native HSA SP and the relevant tag design under the strong pCCW12 promoter as a positive control (**Figure 4A** indicated by black squares). Other samples assayed in this experiment (using Design ɑMF3) covered the high and low extremes of secretion levels, but there was only one mid-range sample present. As such, for the second experiment (using Design ɑMF1), we focused on selecting colonies more in the mid-range to demonstrate the graduated response capacity of the biosensor based secretion assay. Using both tag designs, we showed a robust linear relationship (R^2^ = 0.919/0.934 for tags ɑMF3/ɑMF1, respectively) between biosensor activation and protein secretion. However, tag ɑMF1 had a much shallower slope of ∼2.23×10^-3^, relative to Design ɑMF4 of ∼17.5×10^-3^ (**Figure 4A**). The ɑMF1 tag therefore afforded a much wider dynamic range and sensitivity for the biosensor. Since it also had a minimal effect on protein secretion efficiency, we recommend this design for future work using this system.

From the equation of the dose-response curve (**Figure 1B**), the addition of 5.9 nM of synthetic ɑMF peptide yields a 30-fold increase in fluorescence of the biosensor. This would correspond to a yield of 393 ng/mL for HSA (MW = 66.5 kDa), assuming 1:1 stoichiometry of tag:POI for the ɑMF1 tag. However, this value is not corrected for protein secretion as a function of biomass and the different rates of proteolytic degradation for the secreted tag vs POI over time. We have therefore calculated reciprocal units for both fold-change in fluorescence and protein secretion in ng/ml as a function of culture optical density at 600 nm (OD600, proportional to cell density) when comparing performance of different ɑMF tags (**Figure 3C**). We observed an average maximum cell density of the biosensor strain corresponding to an OD600 value ∼ 9.6 after 72 hours. Using these units of measurements (ng/ml/OD600), we could therefore expect a ∼ 30/9.6 = 3.125 fold increase in GFP Fluorescence/OD600. Using the equation of the line obtained with the ɑMF1 tag (y = 0.002225*x+0.007537), if y = 3.125, x = 1401 ng/ml/OD600, or ∼14 mg/L at a culture of OD600 ∼ 10. For our best secreting ɑMF1 tag construct containing a strong promoter (pCCW12) with the native HSA SP, we observed a maximum of ∼ 0.27 fold increase in Fluorescence/OD600, corresponding to a 115 ng/ml/OD600 yield of HSA. This overall yield agrees with published values for HSA expression in *S. cerevisiae*, albeit with much higher reported OD600 values in the strains used for expression. ^35^ It is important to note that the biosensor strain has been heavily engineered for optimal biosensing and not growth or heterologous protein production. It is likely that yield could be increased dramatically by taking the best performing SP-sequence POI construct identified with the biosensor and using this to express in industrial strains of yeast. One common strategy is via multiple copy integration, in industrial strains of *S. cerevisiae* and potentially other industrially relevant yeasts, such as the methylotrophic *P. pastoris.* ^4, 50^ Shaw *et al.* have demonstrated that the dynamic range of the biosensor is tunable by altering the *LexO* operator repeats within the synthetic promoter controlling reporter gene expression. ^32^ This can be utilised if necessary, to tune the biosensor for other higher expression POI targets.

### Analysis and comparison of optimal protein secretion constructs identified using the ɑMF biosensor from a large a combinatorial library

The first experiment (using Design ɑMF3) identified several expression cassettes with similar secretion efficiency to the native HSA SP control construct, while the second experiment (using Design ɑMF1) focused on identifying clones with intermediate expression (**Figure 4A, Table 1, Supplement File 4**). Table 1 compares the secreted HSA productivity (ng/μL/OD600) of four HSA_ɑMFx4 and 4 HSA_ɑMFx1 expression cassette combinations with the native HSA SP control construct.

**Table 1.**
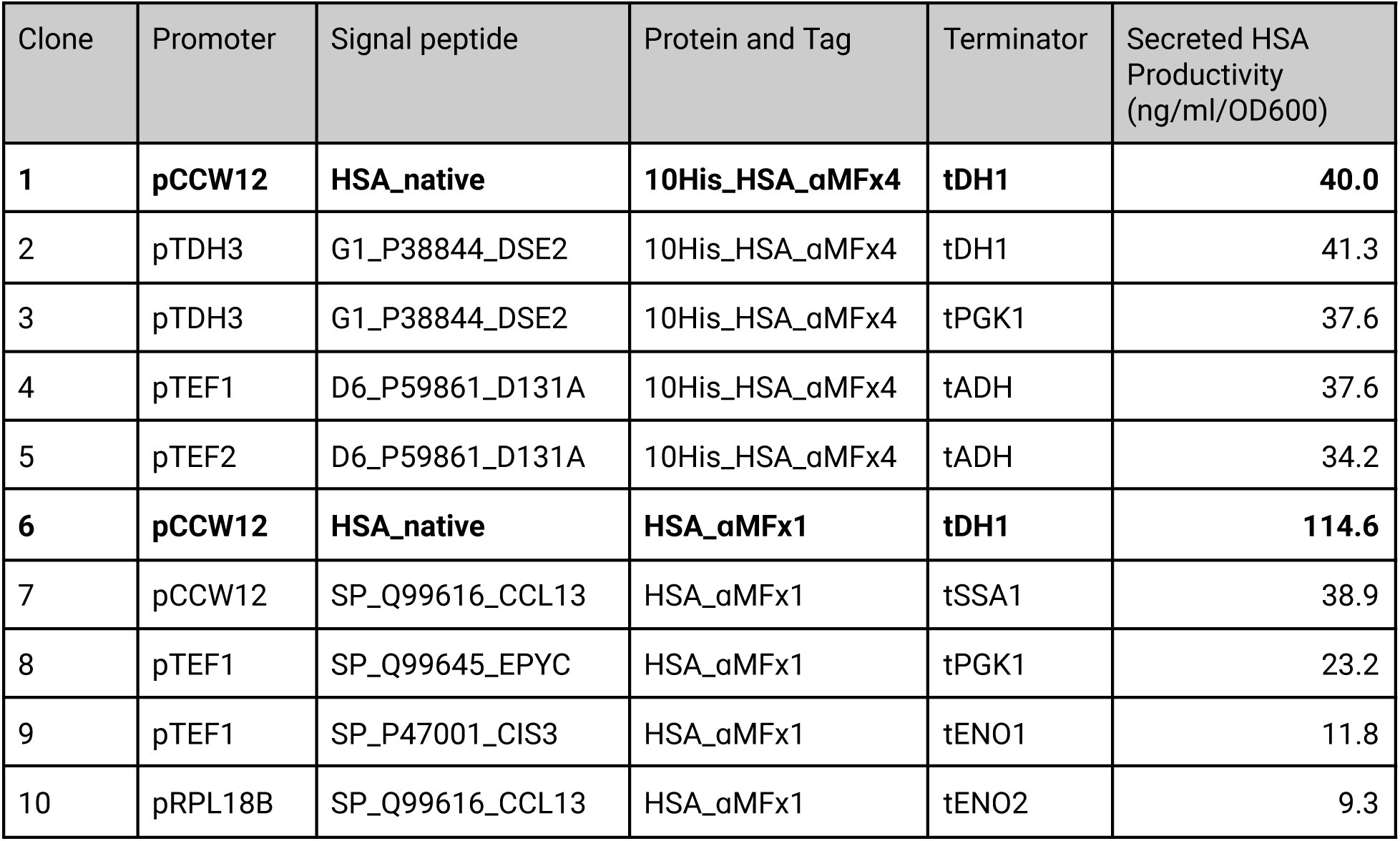
Comparison secreted HSA productivity (ng/mL/OD600) by different HSA_ɑMF expression cassette combinations (for extended table see Supplementary File 4). Positive controls for both ɑMF1 and ɑMF3 tag designs are shown in bold.

| Clone | Promoter | Signal peptide | Protein and Tag | Terminator | Secreted HSA Productivity (ng/ml/OD600) |
| --- | --- | --- | --- | --- | --- |
| **1** | **pCCW12** | **HSA_native** | **10His_HSA_αMFx4** | **tDH1** | **40.0** |
| 2 | pTDH3 | G1_P38844_DSE2 | 10His_HSA_αMFx4 | tDH1 | 41.3 |
| 3 | pTDH3 | G1_P38844_DSE2 | 10His_HSA_αMFx4 | tPGK1 | 37.6 |
| 4 | pTEF1 | D6_P59861_D131A | 10His_HSA_αMFx4 | tADH | 37.6 |
| 5 | pTEF2 | D6_P59861_D131A | 10His_HSA_αMFx4 | tADH | 34.2 |
| **6** | **pCCW12** | **HSA_native** | **HSA_αMFx1** | **tDH1** | **114.6** |
| 7 | pCCW12 | SP_Q99616_CCL13 | HSA_αMFx1 | tSSA1 | 38.9 |
| 8 | pTEF1 | SP_Q99645_EPYC | HSA_αMFx1 | tPGK1 | 23.2 |
| 9 | pTEF1 | SP_P47001_CIS3 | HSA_αMFx1 | tENO1 | 11.8 |
| 10 | pRPL18B | SP_Q99616_CCL13 | HSA_αMFx1 | tENO2 | 9.3 |

The results from the HSA_ɑMFx4 constructs show that our combinatorial library assembly and biosensor can identify novel SPs and constructs with comparable efficiency to the native sequence. Table 1 shows 4 of the highest secreting clones selected,with a selection bias for strong promoters from the MoClo YTK. ^36^ Comparing two constructs (Clones 2 & 3) that differ only in the choice of terminator (tDH1 vs tPGK1) indicates that this part has negligible effects of secretion efficiency in this instance (∼41 vs ∼38 ng/ml/OD600). It is notable that both of these clones by chance were found to contain the identical peptide sequence (G1_P38844_DSE2), evidence of this variable having a strong effect on the selected phenotype.

While the samples tested in the Design ɑMF1 experiment were intentionally selected to be lower than the the native HSA SP control construct, we also observed numerous clones with approximately equal biosensor response to the positive control in this assembly. Furthermore, there were several other insights that were gained from analysing the selected colonies. By selecting for samples with a graduated biosensor response, a broader selection of promoter strengths were observed. However, the strong effect of signal peptide sequence was again illustrated by comparison of clone 7 with the positive control (clone 6), which although contained the strong pCCW12 promoter lacked the the native HSA SP, and displayed a ∼3 fold reduction in secretion efficiency (∼114 vs ∼39 ng/ml/OD600). Moreover, clone 10 possessing the same SP but with the medium strength promoter (pRPL18B) had a further reduction in yield (∼9 ng/ml/OD600). As a further illustration, clones 8 and 9 contained the same medium-strong promoter, pTEF1, but had a 2 fold difference in secretion efficiency based on a different SP. Although all of these examples also had different terminators, the comparison of clones 2 and 3 above, suggests this would not have a strong effect.

Across the two experiments it was clear that the presence of the 10xHis N-terminal tag negatively affected secretion of HSA (**Figure 3C, Figure 4A, Supplementary File 4**). This is likely due to the sequence change around the signal peptidase cleavage site affecting cleavage efficiency. ^16–18^ As shown in **Figure 3C** and comparing the positive controls from both ɑMF tag designs in Table 1 (clones 1 and 6), altering the C-terminal sequence can dramatically alter the secretion efficiency, with longer, more complex tags correlating with less efficient secretion.

Overall, we have presented an efficient method for combinatorial optimisation as applied to heterologous protein secretion in yeast. ^39^ Our system allows identification of the best expression cassette combinations that would otherwise be unlikely to be found, by optimising each component of a genetic construct in isolation.

### Comparison of ɑMF biosensor based protein secretion assay to current methods

Despite being core to the efficient production of industrially relevant proteins, high-throughput methods to optimise the signal peptide sequence for a given POI are not well documented in the literature. Notable exceptions are the secretion of enzymes with fluorogenic or chromogenic substrates, such as ɑ-amylase and β-galactosidase, respectively. These enzymes and substrates have enabled high-throughput screens of signal peptides and genome engineering of strains to improve secretion efficiency, but are limited to optimising these subset of enzymes. ^26, 29, 30^ For the optimization of a more general POI, an invertase-based signal sequence trap method has been used to provide a survival based selection pressure for identifying optimal signal sequences within the genome. ^9, 31^ A potential limitation of this method however, is the requirement for a ∼60 kDa invertase C-terminal tag on a POI, which may limit secretion efficiency. As our results show (**Figure 3C**), even small additions to the C-terminal sequence of a POI can dramatically affect protein secretion efficiency. This invertase-based system also provides only a binary outcome of survival or death, making relative secretion efficiency difficult to quantify. Another useful method is the “*Pichia* secrete and detect” system, that enables rapid generation of SP expression cassettes that can be screened via a split-GFP tag, correlating supernatant fluorescence to secretion yield. ^4, 8^ This method only requires a 16 amino acid GFP11 C-terminal tag, but also requires subsequent addition of a separately purified GFP fragment, and 1-3 nights incubation before reading cultures in a microplate, limiting the throughput and library diversity amenable by this method. ^8^

## Conclusion

In this paper we have presented a novel GPCR biosensor-based protein secretion assay, for use in high-throughput optimisation of recombinant protein secretion from *S. cerevisiae*. We successfully generated and quantified the assembly of thousands of protein expression cassettes from one-pot Combinatorial Golden gate assembly reactions. A limitation of the study is that the protein secretion yields of the biosensor strain are ill-suited to industrial protein production. However, we show that our secretion assay can be used to improve current capabilities in efficiently identifying optimal secretion cassettes for both new and existing POIs. Future work will investigate our hypothesis that these gains in expression efficiency will be transferable when used in industrially relevant yeast strains. In addition, this study tested a limited number of co-translational SPs. Now that the evenness and completeness of one-pot combinatorial assembly using this subset has been verified, this library can be expanded to by including other known SP sequences from *Eukaryota*, including post-translational SPs. Overall, our system provides a rigorous way to analyse the effect of signal peptide sequence on protein secretion efficiency while varying promoter and terminator combinations in parallel, making it a useful tool in furthering our basic understanding of the synergistic effects of these different genetic elements upon this fundamental biological process. This system should also be applicable to high-throughput strain engineering, ^30, 51^ and be useful in further parameterising genome-scale metabolic models of protein secretion.^52^

## Acknowledgments

The authors would like to acknowledge Tom Ellis, William Shaw and colleagues at Imperial College London, for early sharing of the yWS677 engineered strain of S. cerevisiae. The Yeast GPCR-sensor Toolkit was a gift from Tom Ellis (Addgene kit #1000000157). The MoClo-YTK plasmid kit was a gift from John Dueber (Addgene kit # 1000000061). The authors would also like to acknowledge the assistance of the ANU flow cytometry facility and helpful discussion with Elle Saber from the ANU Biological Data Sciences Institute, for helpful discussions regarding statistical analysis of DNA sequencing results. A.W. and J.B would like to acknowledge research funding provided by Samsara Eco and all authors would like to acknowledge additional funding from the Research School of Biology, ANU. A.W., O.B.S and J.B. are members of the ARC Centre of Excellence in Synthetic Biology and the ANU Synthetic Biology Initiative and would like to acknowledge interactions with members of these robust research communities in helping to formulate this research.

## Author contributions

Conceptualization, A.W. and J.B.; Methodology, A.W., R. L., O. B.S, B.S. and J.B.; Software: R.L. and B.S.; Validation, A.W. and J.B.; Investigation, A.W., R. L., O.B.S, L.M. and J.B.; Writing – Original Draft, A.W., R.L and J.B; Writing –Review & Editing, A.W., B.S. and J.B.; Visualization, A.W., R. L., O.B.S and J.B.; Funding Acquisition, J.B.; Project Administration, A.W. and J.B.; Supervision, B.S. and J.B.

## Methods

### KEY RESOURCES TABLE

| REAGENT or RESOURCE | SOURCE | IDENTIFIER |
| --- | --- | --- |
| **Antibodies** |  |  |
| Mouse Monoclonal Human Serum Albumin antibody | Invitrogen by ThermoFisher Scientific | Cat#MA1-19174 |
| Goat Anti-Mouse IgG (H + L)-HRP Conjugate | Bio-Rad | Cat#1706516 |
| Goat Anti-Mouse IgG (H + L)-HRP Conjugate | Bio-Rad | Cat#1706516 |
| **Bacterial and Virus Strains** |  |  |
| NEB<sup>®</sup> Stable Competent *E. coli* (High Efficiency) | New England Biolabs | Cat#C3040H |
| NEB<sup>®</sup> 10-beta Electrocompetent *E. coli* | New England Biolabs | Cat#C3020K |
| **Chemicals, Peptides, and Recombinant Proteins** |  |  |
| α-factor (WHWLQLKPGQPMY) Custom Synthesis (Purity >95%) | Genscript | N/A |
| **Critical Commercial Assays** |  |  |
| Albumin Human ELISA Kit (Sandwich ELISA Kit) | Invitrogen by ThermoFisher Scientific | Cat#EHALB |
| Pierce<sup>™</sup> Bradford Plus Protein Assay Kit | ThermoFisher Scientific | Cat#23236 |
| **Deposited Data** |  |  |
| PlsChain and raw sequencing data | This paper | [10.5281/zenodo.10980037](https://zenodo.org/record/10980037) |
| **Experimental Models: Organisms/Strains** |  |  |
| yWS677 (*S. cerevisiae*, BY4741 derivative) *sst2Δ0 far1Δ0 bar1Δ0 ste2Δ0 ste12Δ0 gpa1Δ0 ste3Δ0 mf(alpha)1Δ0 mf(alpha)2Δ0 mfa1Δ0 mfa2Δ0 gpr1Δ0 gpa2Δ0* | Shaw et al. 2019 | N/A |
| αMF protein secretion biosensor strain (*S. cerevisiae*, yWS677 derivative) pRAD27-LexAPRD-tEN01, pLexOx6-sfGFP-tTDH1, pPGK1-Gpa1-tTENO2, pCCW12-Ste2-tSSA1. | This paper | N/A |
| **Oligonucleotides** |  |  |
| For oligonucleotides used as primers and for creating gRNAs see Supplementary file 4 | This paper | N/A |
| **Recombinant DNA** |  |  |
| Yeast MoClo Toolkit | Lee et al. 2015 | Addgene Kit: 1000000061 |
| Yeast GPCR-sensor Toolkit | Shaw et al. 2019 | Addgene Kit: 1000000157 |
| 46 Signal peptides<br>(Full sequences available in Supplementary file 3 and Supplement Benchling Sequences) | Twist Biosciences | Custom Synthesis, N/A |
| **Software and Algorithms** |  |  |
| Prism | GraphPad | https://www.graphpad.com/scientific-software/prism/<br>RRID: SCR_002798 |
| Benchling | Benchling | https://www.benchling.com<br>RRID: SCR_013955 |
| UniProt | The Universal Protein Resource | http://www.uniprot.org/<br>RRID:SCR_002380 |
| PlsChain | This paper | N/A |

#### Resource availability

Further information and requests for resources and reagents should be directed to and will be fulfilled by the lead contact

#### Materials availability

Plasmids generated in this study are in the process of being deposited with Addgene.

#### Data and code availability

Data have been deposited at Zenodo (https://zenodo.org/doi/10.5281/zenodo.10980037) and are publicly available as of the date of publication. Accession numbers are listed in the key resources table. All raw sequencing data analysed with PlsChain is available at Zenodo. DOIs are listed in the key resources table. Code for PlsChain is publicly available at GitHub https://github.com/RunpengLuo/PlsChain. Other scripts used in this research are available in Supplementary file 3.

Any additional information required to reanalyze the data reported in this paper is available from the lead contact upon request.

#### Experimental model

##### Yeast strains

The protein secretion biosensor *S. cerevisiae* strain used in this study is derived from yWS677 described in Shaw *et al*., ^32^ which is derived from the stable haploid strain BY471, ATCC: 4040002 (*MATα SUC2 gal2 mal2 mel flo1 flo8-1 hap1 ho bio1 bio6 is3Δ1 leu2Δ0 met15Δ0 ura3Δ0*). See key resource table above.

### Method details

#### Bacterial Strains and Growth Media

NEB® Stable Competent *E. coli* (High Efficiency) was used for general cloning experiments. NEB^®^ 10-beta Electrocompetent *E. coli* was used for Combinatorial Golden Gate *E. coli* transformation experiments. Growth and selection of *E. coli* was performed in Lysogeny Broth (LB) medium at 37°C with aeration. LB medium was supplemented with appropriate antibiotics for selection at each stage of cloning (ampicillin 100 μg/mL, chloramphenicol 34 μg/mL, or kanamycin 50 μg/mL).

#### Yeast transformations

Yeast transformations were performed based on the lithium acetate protocol.^53^ Full protocol available on [Benchling](https://benchling.com/x_josbr/f_/wrOmoFXE-protein_secretion_biosensor_ms_2024/).

#### Bacterial transformations

Bacterial transformations followed either High Efficiency Transformation Protocol using NEB® 10-beta Competent *E. coli* (High Efficiency) (#C3019H, NEB) or Electroporation Protocol (#C3020, NEB). For electroporation, exponential protocol setting (Voltage 1800V; Capacitance 25 uF; Resistance 200 ohms; Cuvette 1mm) was used on GenePulser Xcell (BioRad).

#### Yeast Strains and Growth Media

For details of yeast strains used in this study see Key Resources Table. The αMF biosensor strain (*S. cerevisiae,* yWS677 derivative) *STE2 GPA1 LexA-PRD sfGFP* was generated using markerless CRISPR/Cas9 genome engineering (see **Construction of the Biosensor Strain**, below).

Yeast extract peptone dextrose (YPD) was used for culturing cells prior to transformation: 1% (w/v) Bacto Yeast Extract (Sigma), 2% (w/v) Bacto Peptone (Sigma), 2% glucose (Astral). Cells were cultured at 30°C shaking at 250 rpm.

Selection of yeast transformants was performed on synthetic complete (SC) uracil dropout agar medium: 2% (w/v) glucose (Astral), 0.67% (w/v) Yeast Nitrogen Base without amino acids (Sigma), 0.14% (w/v) Yeast Synthetic Drop-out Medium without uracil without nitrogen base (USBiological) supplemented with 2% (w/v) bacteriological agar (Astral). For experiments selecting against the *URA3* auxotrophic marker following CRISPR/Cas9 genome engineering, SC-URA dropout media was supplemented with 20 mg/L uracil (Astral) and 0.1% (w/v) 5-Fluoroorotic acid (5-FOA). Cells were grown at 30°C static.

All liquid experiments were performed in synthetic complete uracil dropout (-URA) medium: 2% (w/v) glucose (Astral), 0.67% (w/v) Yeast Nitrogen Base without amino acids (Sigma), 0.14% (w/v) Yeast Synthetic Drop-out Medium without uracil without nitrogen base (USBiological). For 72 hr protein expression experiments the medium was supplemented with 100mM Potassium phosphate buffer (pH 6.5) Unless otherwise stated, all yeast strains were cultured in 10 mL of SC-URA medium and grown in 50 mL falcon tubes at 30°C, with orbital shaking at 250 rpm.

#### Golden Gate assembly

Golden Gate assembly reaction mixtures were made in 0.2 mL PCR tubes as follows:

**Table M1.**
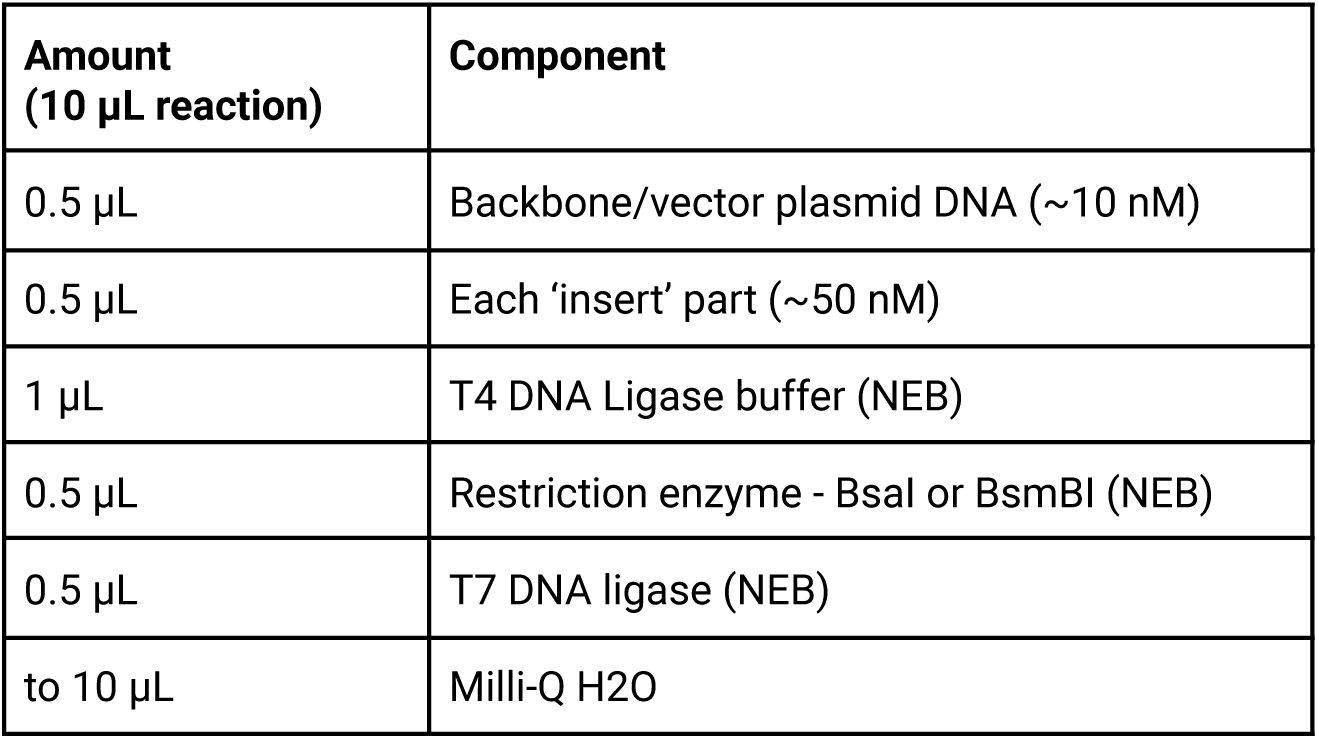
Standard Golden Gate reaction mixture.

All Golden Gate assembly reactions were performed using a thermocycler (Biometra T300) with
the following program:

**Table M2.** Golden Gate assembly Thermocycler incubation program.

| Step | Temperature | Time |
| --- | --- | --- |
| 25 cycles of digestion and ligation | 42 °C | 2 minutes |
|  | 16 °C | 5 minutes |
| Final digestion or ligation step * | 60 °C or 42 °C | 10 minutes |
| Heat inactivation step | 80 °C | 10 minutes |
| Hold | 4°C | - |

*Final ligation step (42 °C) was performed when assembling plasmids to be used as integration or expression vectors. Final digestion step (60 °C) was used for all other assemblies. The final digestion step (60 °C) reduces the level of intact vector background.

#### DNA Extraction Protocols

All plasmid *E.coli* plasmid DNA minipreps were performed using the NEB Monarch Plasmid Miniprep Kit and Protocol (NEB #T1010). Prior to starting the miniprep protocol, 5 mL overnight cultures were pelleted using a 5810-R centrifuge (Eppendorf) at 800 x *g* for 5 minutes. Supernatant was removed. Some cultures were stored at −20 °C at this point for several days before being defrosted slowly on ice. Fresh or defrosted *E. coli* pellets were gently resuspended in 500 μL of sterile Milli-Q H2O and transferred into a sterile 1.5 mL microcentrifuge tube. The Monarch Plasmid Miniprep Protocol (NEB #T1010) was followed from here with no known deviations. A Heraeus Pico 21 Benchtop microcentrifuge (ThermoFisher Scientific) was used for all centrifugation steps. An optional step of heating elution buffer to 50 °C prior to elution, to improve yield, was followed. Final elution volume for all preps was 50 μL.

Yeast genomic DNA was extracted from the potential aMF biosensor yeast strains and the unedited yWS677 strain (to be used as negative control in PCR analysis) using the YeaStar Genomic DNA kit and protocol (Zymo Research), with no known deviations.

Yeast plasmid DNA extractions were performed by preparing overnight liquid cultures from single yeast colonies, plasmid DNA was extracted via the Yeast Miniprep kit II and protocol (Zymo Research), with no known deviations.

#### Construction of the Biosensor Strain

The biosensor gene constructs were designed using the Benchling Virtual Golden Gate Assembly tool **(See figure S3 for plasmid map and** [Benchling project](https://benchling.com/x_josbr/f_/wrOmoFXE-protein_secretion_biosensor_ms_2024/)). Following successful virtual assembly, the actual constructs were assembled. The Level 1 gene assemblies for the biosensor receptor (Ste2), G-protein alpha-subunit (Gpa1), synthetic transcription (LexA-PRD) factor and reporter gene (sfGFP) were assembled via Golden Gate assembly using Level 0 parts from the MoClo YTK and Yeast GPCR-sensor Toolkit. ^32, 36^ A markerless integration vector with ‘234’ GFP dropout site was also assembled using Level 0 parts from these kits. The four Level 1 constructs were assembled, via Golden Gate assembly, into the markerless integration vector to create a Level 2 construct. Following sequence confirmation of the Level 2 integration cassette containing the biosensor genes, the genes were inserted into the host yeast genome via CRISPR/Cas9 mediated gene editing. The integration cassette contained 500 bp homology arms compatible with the LEU2 landing pad site present in the yWS677 yeast strain genome. Linearised single guide RNA (targeting the LEU2 landing pad site), a linearised CRISPR/Cas9 sgRNA expression vector (containing *URA3* auxotrophic marker gene) and linearised Level 2 biosensor gene construct (donor DNA) was transformed into the yWS677 yeast strain. The guide RNA and linearised CRISPR/Cas9 sgRNA expression vector have compatible homology arms for use in gap repair once in the yeast cell. Cells proficient in gap repair were selected for by spreading the transformants onto-URA 2% glucose 2% agar plates. Successful transformants were then screened for successful genomic integration of the biosensor genes via a series of PCRs (for guide RNA and PCR primer sequences see **Supplementary file 4**). Following confirmation of successful gene integration the resulting biosensor strain was cured of any remaining CRISPR/Cas9 sgRNA expression vector via selection against the *URA3* auxotrophic marker gene using 5-Fluoroorotic acid (5-FOA). Plasmid ‘curing’ was performed by growing a liquid culture of the biosensor strain in YPD (a non-selective media) overnight (30 °C, shaking at 200 rpm). The culture was passaged several times in liquid YPD media, to allow time for most of the yeast cells to lose the Cas9 sgRNA plasmid (containing the URA3 gene). Cells were then plated onto 5-FOA Synthetic Complete (SC) media agar plates and incubated for 72 hours at 30 °C. A single colony was selected and streaked onto a fresh 5-FOA SC media agar plate and again incubated for 72 hours at 30 °C. A single colony was selected from the new plate and use to inoculate 10 mL of liquid YPD (grown overnight, 30 °C, shaking at 200 rpm). The overnight culture was spotted onto both a YPD agar plate and a-URA 2% glucose 2% agar plate, to confirm that the biosensor yeast could no longer grow on selective media lacking Uracil.

### Testing activation of *S. cerevisiae* ɑMF biosensor yeast strain

The potential ɑMF biosensor strain was screened for biosensor activation by applying water (-ve control) or 100 nM synthetic *S. cerevisiae* ɑMF peptide (**Figure S3D**). Post hoc reverse pairwise comparisons and a Two-way ANOVA were performed in R to analyse these results (See **Supplementary file 3**).

### Construction of Dose-response curve

In a sterile 2.2 mL 96 deep-well plate, a single colony of the biosensor strain was inoculated into 500 μL of SC liquid media. The liquid culture was left to grow to stationary phase overnight at 30 °C, shaking at 200 rpm. The culture was back diluted 1/100 in SC liquid media into 10 wells of the deep-well plate. The plate was left to grow for 1 hr at 30 °C, shaking at 200 rpm, then varying concentrations of synthetic *S. cerevisiae* aMF peptide were added to each well (0, 0.01 nM, 0.1 nM, 0.316 nM, 1 nM, 3.16 nM, 10 nM, 31.6 nM, 100 nM, 1 μM). The plate was left to grow for a further 4 hrs at 30 °C, shaking at 200 rpm, before results were recorded using a BDTM LSRII cytometer (MCRF) with GFP parameter voltage set to 583 V. Single cell populations were determined via FSC-A, FSC-H and FSC-W (parameter voltage set to 525 V). Only single cells were selected for fluorescence analysis. 20,000 single cells were analysed for each sample.

The dose-response curve was plotted using Prism 9 (GraphPad). The raw results were transformed to a log10 scale (Analyse → Transform concentration (X) → Log10 transformation) and plotted on an XY scatter plot. Four parameter non-linear regression (curve fit) of transformed results was performed using Prism 9 (GraphPad) (Analyse→XY analysis → Non-linear regression (curve fit) → log(agonist) vs. response – Variable slope (four parameters)). Values obtained from the ‘Nonlin Fit’ table of results in Prism 9 (GraphPad), for the four-parameter non-linear regression (curve fit), were used to assess the goodness of fit of the dose-response curve as shown in the equation below:

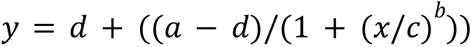

The span of the curve is 35.21 units (fold over background). The four parameters of the ‘best fit’ dose-response curve, as calculated in Prism 9 (Graph pad), are Bottom (a) = -1.448, Top (d) = 33.77, EC50 (c) = 7.869e-10, HillSlope (b) = 1.052. The equation of the ‘best fit’ dose-response curve is:

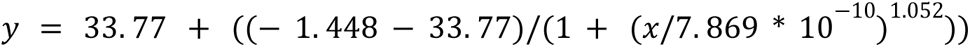

Where *x* is the dependant variable (Dose, ɑMF (Moles/L)) and *y* is the independent variable (Response, GFP fluorescence fold above background).

### SP dataset collection

Amino acid sequences of all characterised secretory proteins of a particular domain, genus or species were obtained from UniProt (The UniProt Consortium, 2020). Advanced search was used, with the input “locations:(location:“Secreted [SL-0243]”) annotation:(type:signal) AND reviewed:yes”. Separate sets of sequences were collected for various domains (AND taxonomy: 2759), genera (AND taxonomy: 4930) and organisms (AND organism: 5400, 105351, 5061, 38033, 5346, 685588, 9606, 4911, 4922, 573729, 870730, 4932, 4952). Sets of sequences obtained from UniProt in the FASTA format were submitted to SignalP 5.0 (organism group: Eukarya, output format: long format). The BEDTools suite (bedtools v2.30.0) was used to extract the FASTA of processed SPs from the processed entries GFF3. Additional amino acids in the native protein sequence were appended to the sets of SP sequences after the cleavage site calculated by SignalP 5.0 using AWK.

### SP diversity analysis and selection

Sequences were aligned using the MAAFT online service. Compute Pairwise Distance in MEGA11 was used to calculate distance matrices using the Dayhoff model. Truncated single value decomposition (SVD) followed by t-distributed stochastic neighbour embedding (t-SNE) were applied to each distance matrix. Both operations were implemented in scikit-learn. Diverse SPs were visually identified on the three-dimensional scatter plot based on maximising Euclidean distance within the plot to other points. Sequences were subsequently aligned using Clustal Omega (**Supplementary file 3**).

### Design of the 3a-3b AP cloning junction

Due to the requirement for Golden Gate cloning junction within open reading frames to be standardised, we needed to make a choice to redefine the MoClo YTK assembly standard^36^ of a glycine-serine linker at the type 3a-3b junction to ensure the best possible signal peptide cleavage efficiency for all sequence combinations. Our analysis using Sequence Logo^44^ **(Figure 2B**) showed a qualitative enrichment for alanine and proline at positions +1 and +2, respectively, within Eukaryotic signal peptide sequences. After considering all possible codons to achieve this substitution in comparison with other MoClo YTK overhangs and *S. cerevisiae* codon usage preferences, we found that the type8-type1 junction (CCCT) defined within the MoClo YTK assembly standard would be most suitable **(Figure 2C**). This would use the Ala-Pro codons: GCC CCT. Although already in use within the assembly standard, type 8 (Bacterial marker and origin) and type 1 (assembly connectors) are generally used when constructing plasmid backbones. The overhang can be re-used if expression or integration plasmids are first constructed as ‘234-dropout’ plasmids, as is common practice and described in the original publication. ^36^ Users of our system should be aware of this necessity to first construct a drop-out vector backbone when designing expression constructs using our signal peptide library.

### Combinatorial Golden Gate assembly of expression cassette plasmid libraries

All expression constructs were assembled into a yeast episomal expression vector (YEE) with a type ‘234’ *mRFP* dropout site. This allowed for red/white screening of *E. coli* colonies for successful Golden Gate assembly, followed by restriction digest analysis and whole plasmid sequencing to verify correct assembly of individual constructs. Combinatorial Golden Gate assembly was performed by preparing equimolar concentration Level 0 part plasmid libraries/ mixtures. A sample of each Level 0 plasmid part was prepared, then a library master mixture containing equal equimolar was prepared for each; SPs, promoters and terminators. Combinatorial Golden Gate reaction mixtures were prepared as in **Table M1**, with the Level 0 part plasmid library master mixtures substituted for single ‘insert’ parts. The reaction conditions were the same as for Golden Gate assembly (**Table M2**).

Following assembly the ligation mixture was transformed via electroporation into NEB^®^ 10-beta Electrocompetent *E. coli*, according to the NEB Electroporation Protocol (#C3020, NEB). Transformants were plated onto LB-Ampicillin 2% agar media in large (150mm) petri dishes. A ‘batch miniprep’ plasmid DNA extraction was performed for each transformation.

Transformant colonies (>3x expected library diversity) from a single transformation were selected from the plates and resuspended in 15 mL liquid LB containing ampicillin. A sterile plastic cell spreader was used to gently loosen the colonies on the surface of the agar. 2 mL of liquid LB containing ampicillin was pipetted onto the surface of the plate, then the plastic cell spreader was used to gently scrape the colonies from the surface of the plate. The colonies were resuspended in the 2 mL of liquid media and transferred to a 50 mL falcon tube. A second wash of 1-2 mL of liquid LB containing ampicillin was pipetted onto the surface of the plate, then the plastic cell spreader was used to collect any remaining cells. The cells were resuspended in the liquid media and transferred to the 50 mL falcon tube, where liquid LB containing ampicillin was added to a final volume of 15 mL. Depending on the transformation efficiency of the sample, 3-5 plates of colonies were needed to reach >3x expected combinatorial library diversity. The 15 mL cell solution was mixed well to ensure full resuspension of cells and to eliminate any cell clumping. 1.5 mL aliquots were then taken for plasmid DNA extraction. Plasmid DNA miniprep was performed and DNA concentrations were measured as above (see Method *E. coli* DNA extraction protocol). The resulting extracted plasmid DNA library was sequenced via whole plasmid Oxford Nanopore sequencing and analysed using the PlsChain algorithm.

### Screening expression cassette libraries for improved protein secretion via a yeast based biosensor

Combinatorial POI expression cassette plasmid libraries were transformed into the ɑMF biosensor yeast strain for screening. The resulting transformants were plated out on selective media at an appropriate dilution to ensure that the majority of colonies were singular.

Plates of transformants were imaged using a Biorad ChemiDoc Touch Imaging System (Bio-Rad) with black background, auto-exposure, on the Alexa488 (blue epi light) and Colorimetric (white light) settings (located in the ‘Blots’ menu). Alexa488 images were manually contrast adjusted until only the fluorescent colonies could be seen. The images were then overlaid and false coloured (Alexa488 in green, Colorimetric in red) to allow for visualisation of green-fluorescent vs non-fluorescent colonies. Image J software was used to count the total number of colonies and to identify the brightest colonies via the ‘Find Maxima’ tool. Following initial analysis in Image J, Image Lab software was used to quantify the maximum RFU of chosen colonies. Selected colonies were then grown as liquid cultures. Flow cytometry was used to accurately quantify single cell GFP fluorescence of these cultures, and yeast plasmid DNA was extracted via plasmid DNA miniprep using Yeast Miniprep kit II (Zymo Research) then sequenced via whole plasmid Oxford nanopore sequencing to determine the successful expression cassette combinations.

Plasmid sequencing results were analysed using the Benchling sequence alignment tool. The consensus sequence FASTA files were imported as DNA sequences in Benchling, then the sequence alignment tool was used to compare all possible components for each of the part libraries (promoters, signal peptides and terminators). Based on which parts aligned best, *in silico* Golden Gate assembly was performed to re-create the plasmid in Benchling. This plasmid map was then re-compared to the sequencing results to confirm absence of any mutations in the total expression cassette and vector.

Total secreted HSA concentrations of 72 hr yeast HSA expression cultures were quantified via commercially available Albumin Human ELISA Kit (ThermoFisher Scientific #EHALB), following the included protocol. Cultures were inoculated from single colonies containing different HSA expression cassettes, then grown overnight. The optical density of the overnight cultures was read and new cultures were set up via back-dilution to standardise samples to OD600 = 0.2 (pre-log growth phase). The resulting cultures were grown for 72 hours, then harvested for analysis. Flow cytometry was used to accurately quantify single cell GFP fluorescence of these cultures, optical density (OD600) was quantified via spectrophotometry and an aliquot of the remaining culture was centrifuged to pellet the cells. The culture supernatant was used for HSA ELISA analysis. Samples that were expected to have high levels of HSA secretion were diluted 1:1 with fresh -URA 2% glucose media, prior to running the ELISA protocol; to ensure all samples were within the range of the ELISA HSA standard curve. All samples and standards were run in triplicate.

### SDS-PAGE and Western Blot analysis

50 mL liquid cultures of each sample were grown in 250 mL conical flasks, covered with foil (shaking at 200 rpm, 30°C, 72 hrs). The cultures were set up by back diluting overnight starter cultures to OD600 = 0.2 in potassium phosphate buffered -URA 2% glucose liquid growth medium. After 72 hrs of growth, the cultures were harvested. OD600 of each culture was measured via SpectraMax QuickDrop Micro-Volume Spectrophotometer (Molecular Devices LLC). A 1 mL aliquot of each expression culture were pelleted using a Heraeus Pico 21 Benchtop microcentrifuge (ThermoFisher Scientific) (4000 x g, 2 minutes), and supernatant was removed (cell pellet to be used for total cellular protein analysis). The remainder of the expression cultures were pelleted using a 5810-R centrifuge (Eppendorf) (800 x g, 10 minutes) and the supernatant was collected (for secreted protein analysis). 30 mL of each supernatant sample was concentrated 100-fold using an Ultra-15 30K (Amicon) centrifugal filter unit to obtain concentrated secreted protein. *S. cerevisiae* cell pellets were resuspended in 0.5 mL ice-cold EME extraction buffer, then lysed via passage through a French pressure cell (750 MPa) to obtain total cellular protein (cell lysate). 50 μL of cell lysate was centrifuged (21,000 x g, 2 minutes, 4 °C) to obtain soluble cellular protein (in supernatant).

A Bradford assay was used to determine total protein concentrations of the samples. Protein concentrations of the soluble cellular protein and 100x concentrated secreted protein samples were quantified against Bovine Serum Albumin (BSA) standards using the Pierce Coomassie Protein Assay kit (#23236, ThermoFisher Scientific). The assays were performed in 96-well transparent plates containing 150 μl of diluted protein sample or 0, 0.75, 1.5 and 2.25 μg BSA (used to generate the standard protein curve) to which 150 μl of Bradford Coomassie reagent was added. After 10 min at room temperature the absorbance at 595nm of each well was measured using a xMarkTM Microplate Absorbance Spectrophotometer (Bio-Rad). Results of the assay were used to standardise the amounts of cell lysate sample loaded for SDS-PAGE.

Cell lysate, concentrated secreted protein and HSA protein standard stocks were diluted 1:1 with 2X SDS buffer. The samples were boiled for 5 minutes and centrifuged using a Heraeus Pico 21 Benchtop microcentrifuge (ThermoFisher Scientific) (14,000 x g for 5 minutes), before loading onto pre-cast 4-12% Bis-Tris buffered NuPAGE gels (Invitrogen) along with a Precise Plus protein standard ladder (Bio-Rad). 14 μL of each cell lysate sample (1.3 μg of soluble protein, plus insoluble/membrane proteins) and 10.9 μL of each concentrated (50x) supernatant sample was loaded for both gels, one for Coomassie staining and one for Western blotting. For the gel intended for Coomassie staining 0.5 ng and 0.25 ng of HSA protein standard was loaded. For the gel intended for Western blot analysis 0.05 ng, 0.025 ng and 0.0125 ng of HSA protein standard was loaded.

PAGE separated proteins were transferred onto a PVDF membrane (AmershamTM Hybond®) in transfer buffer using a XCell IITM Blot Module (Invitrogen) at 25 V, 4 °C for 70 minutes. Following transfer, the membrane was incubated in blocking reagent for 1 hour at room temperature with gentle orbital agitation. The membrane was washed with TBS, then incubated overnight at 4 °C with primary antibody (mouse Monoclonal Human Serum Albumin antibody #MA1-19174, invitrogen) diluted 1:4000 in TBS buffer. The membrane was washed with TBS buffer (3 times for 10 minutes each) to remove all unbound antibody. The membrane was then incubated for 1 hour with a secondary antibody (Horseradish peroxidase (HRP) conjugated goat-anti-mouse antisera, BioRad) diluted 1:2000 with TBS. The membrane was washed with TBS (3 times for 5 minutes each), then incubated with ECL clarity substrate (Bio-Rad) for 30 seconds. Immunoreactive bands were immediately imaged using a ChemiDoc XRS+ System (Bio-Rad), ‘Chemiluminescence’ setting, exposure time 50 seconds.

### Creating a combinatorial library analysis program

Classification of sequencing reads onto the library of expression cassettes can assist downstream analysis on accessing the library diversity and completeness. Long read mappers such as Minimap2 can classify the reads by aligning them onto all possible combinations of expression cassettes as references and pick the primary alignments. However, such an approach may suffer from extremely high computational costs with respect to the exponential number of reference sequences, while maximising the alignment sensitivity is unnecessary to solve the classification problem. Hence, we developed PlsChain, a highly efficient and accurate classifier that precisely classifies sequencing reads based on unique *k*-mers and branching.

#### Preliminary

We assume 0-indexing throughout this section. An *ordered set* is a pair (*P,C*) where *P* is a set and *C* is an *order* on *P*. *C* is *linear* if *C* ⊆ *P*^2^, whereas *C* is *cyclic* if *C* ⊆ *P*^3^. *C* is *partial* if not all pairs of elements in *P* are comparable, *total* otherwise. All the ordering mentioned in the following subsections are assumed to be total unless stated explicitly. Given a linearly ordered set (*P,<*), for any *a,b* ∈ *P*, *a* ≠ *b*, we say *a* precedes *b* when *a < b*. (*P*[*i*])_0≤*i<*|*P*|_ is a *sequence* over set *P* arranged by the linear order *<*, such that *P*[0] *<…< P*[|*P*| − 1]. If (*P,C*) is a linearly (cyclic) ordered set and *H* ⊆ *P*, then *C* ∩*H*^2^ (*C* ∩*H*^3^) is an order on *H*, denoted as *C*|*_H_*, and (*H,C*|*_H_*) is an *ordered subset* of the ordered set (*P,C*).

Let (*P,C*) be a cyclic ordered set^54^, a *cut* on (*P,C*) is a linear order *<* on *P*, such that for any *x,y,z* ∈ *P*, *x < y < z* ⇒ (*x,y,z*) ∈ *C*. *<_C,x_* denotes a cut with the least element *x* ∈ *P* on (*P,C*). Similarly, given a linear order *<* on *P*, we can construct a cyclic order *C_<_* on *P* by (*x,y,z*) ∈ *C_<_* if and only if either *x < y < z* or *y < z < x* or *z < x < y* for any *x,y,z* ∈ *P*.

A *polytree P* = (*V,E*) is a directed acyclic graph whose underlying undirected graph is a tree. The vertices without any in-coming edges are defined as *root*, whereas the vertices without any out-going edges are defined as *leaf*. For any edge (*u,v*) ∈ *E*, *u* is the *parent* of *v* and referred to as v.p. A *root-to-leaf* (RTL) path *p_u,v_* is a directed path that begins with a root *u* and ends with a leaf *v*. An *arborescence P_u_* = (*V,E*) is a polytree with exactly one root *u*, such that all edges are directed away from *u*. The *depth* of a vertex *v* ∈ *V* (*P_u_*) is the number of edges in the RTL path *p_u,v_*, where the root vertices have depth as 0. A *branching B* = (*V,E*) is a forest of arborescences, which is a directed graph with all of its connected components are arborescences. *P_B,u_* denotes an arborescence with root *u* in a branching *B*.

#### Problem formulation

The concatenation of the strings from *n* plasmid components in a cycle forms the genome sequence of the plasmid. Let Σ = {*A,C,G,T*} be the alphabet of nucleotides, *S_i_* is the *i*th plasmid component that consists of a set of strings over Σ. Component *S_i_* precedes *S_j_* if (*i* + 1) ≡ *j* (mod *n*). Theoretically, there are 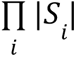 possible plasmid combinations. Given a group of plasmid components, the problem is to classify the sequencing reads against all possible combinations.

#### Indexing

A *k*-mer is a substring of length *k* over alphabet Σ. We define the canonical representation of a *k*-mer as the lexicographically smallest one among itself and its reverse complement representation, and the reverse complement can be computed in *O*(1) using a lookup table. The *k*-mer *identity* (*α,β*) indicates that the *k*-mer is originating from string *s_β_* over plasmid component *S_α_*, with *α* ∈ {0,1*,…,n*−1} and *β* ∈ {0,1*,…,*|*S_α_*| − 1}. The hash function *ϕ* over alphabets is defined as *ϕ*(*A*) = 0, *ϕ*(*C*) = 1, *ϕ*(*T*) = 2, and *ϕ*(*G*) = 3. During the indexing step, the canonical *k*-mers that uniquely present in exactly one genome sequence among all components are hashed via hash function *ϕ* and inserted to a hash table *HT* as the keys, while the values are representing the *k*-mer identities.

#### Classification

A *k*-mer is *informative* if it appears in the pre-built hash table *HT*, *noninformative* otherwise. Given a query string *Q*, we select all the informative *k*-mers from *Q* in a sliding window approach and record their *k*-mer identities by extracting the value from *HT*. An anchor *a* represents an informative *k*-mer, it consists of a quadruple (*α,β,q,c*), such that the *k*-mer is at *q*th position of *Q* with *k*-mer identity (*α,β*), where *k*-mer count *c* = 1.

Let (*A,<_q_*) be a linearly ordered set, *A* is a set of *m* anchors over *Q* and *<_q_* is a linear order on *A*. For any *a,b* ∈ *A*, *a <_q_ b* ⇔ *a.q < b.q*. (*A*[*i*])_0≤*i<m*_ is a sequence of anchors in *A* ordered by *<_q_*. Adjacent anchors sharing the same *k-*mer identity will be merged into one by increasing the *k*-mer count accordingly. In addition, we define the partial order *<_α_*⊆ *A*^2^ on set *A*. For any *a,b* ∈ *A*, *a <_α_ b* ⇔ *S_a.α_* precedes *S_b.α_*. To account for the circular genome structure, we define the ternary relation *C* = *C_<q_* ⊆ *A*^3^, such that (*A,C*) forms a cyclic ordered set.

A *chain* (*H,C*|*_H_*) is a cyclic ordered subset of the cyclic ordered set (*A,C*) with |*H*| = *n*. Let *<* be any cut on (*H,C*|*_H_*), given the sequence (*H*[*i*])_0≤*i<n*_ ordered by *<*, for any integer *j* ∈ [0*,n* − 1), *H*[*j*] *<_α_ H*[*j* + 1]. The weight of (*H,C*|) can be computed by the total *k*-mer count, *i.e*. Σ_{*u∈H*}_ 𝑢. 𝑐. Thus, we can formulate the plasmid classification problem as finding an optimal chain (*H_opt_,C*|*_Hopt_*) with maximum weight.

Notice that there exists 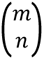 distinct subsets of *A* with size *n*, and test for the partial ordering constraint *<_α_* per combination takes *O*(*n*). Thus, the naive solution would take 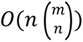 to decide the optimal chain, which is impractical when *n* ≪ *m*. Therefore, we proposed PlsChain, an iterative algorithm to update a branching and finally obtain the optimal chain through RTL path traversal. Given a branching *B* = (*V,E*), every vertex *v* ∈ *V* (*B*) can represent an anchor *a* ∈ *A*, denoted as *v.a*. Let *P_B,u_*be an arborescence with root *u* ∈ *V* (*B*), *u.a* defines a cut *<_C,u.a_* on the cyclic ordered set (*A,C*). Let *v* ∈ *V* (*P_B,u_*), for any vertex *w* ∈ *V* (*B*)\{*v*}, (*v,w*) ∈ *E*(*B*) only if both *v.a <_α_ w.a* and *v.a <_C,u.a_ w.a*. If there exists a vertex *w* ∈ *V* (*B*) that may connects to two distinct vertices 𝑣_1_ ∈ 𝑃_𝐵,𝑢_1__ and 𝑣_2_ ∈ 𝑃_𝐵,𝑢_2__, *w* will be duplicated into two vertices *w*_1_ and *w*_2_ and connected to *v*_1_ and *v*_2_, respectively. Notice that any vertex with depth *n*−1 is a leaf in *B*, which is the maximum depth in *B*. Thus, by progressively inserting the vertices and edges to the branching *B*, we can obtain a set of chains through RTL path traversal against the leaves with depth *n*−1, and select the optimal chain (*H_opt_,C*|*_Hopt_*) accordingly. The *k*-mer identities of the sequence (*H_opt_*[*i*])_0≤*i<*|*Hopt*|_ ordered by *<_α_* forms the class of the query string *Q*. Conversely, query strings that lead to no chains are left unclassified by PlsChain.

In practice, given a plasmid library with *n* plasmid components, an informative *k*-mer ‘survives’ without errors in a read with the survival rate 𝑠𝑢𝑟𝑣𝑖𝑣𝑎𝑙𝑅𝑎𝑡𝑒(𝑘) = (1 − 𝑝)*^k^*, where 𝑝 is the probability of an error at a given position of a read with respect to varying sequencing technologies. ^55^ A read is classified if it contains at least *n* informative *k*-mers that coming from *n* different plasmid components, which leads to an approximated classification rate 𝑐𝑙𝑎𝑠𝑠𝑖𝑓𝑖𝑐𝑎𝑡𝑖𝑜𝑛𝑅𝑎𝑡𝑒(𝑘) = (1 − 𝑝)^𝑘𝑛^. With p=0.05, k=15, and n=6, the theoretical classification rate is as low as 9.88%. Thus, we introduced *partial-classification* to rescue error-prone (but informative) reads. A read can be partially classified if there exist informative *k*-mers against some but not all plasmid components. During the process of updating the branching, PlsChain inserts artificial vertices and edges to the branching if there does not exist any informative *k*-mers for the corresponding plasmid components at certain depth. Artificial vertex inherits the anchor from its parent vertex with *k*-mer identity marked as asterisk. If a read only misses informative *k*-mers coming from plasmid components with a single sequence of choice, PlsChain can even fuzzy match the read by inferring the missed plasmid component accordingly. Hence, PlsChain is capable of partially classifying error-prone reads to several potential plasmid combinations instead of leaving them unclassified. In addition, PlsChain pre-filters reads potentially sourcing from contamination during sequencing. Given the theoretical maximum sequence length of a plasmid sequence over the library of n expression cassettes is 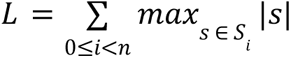. For any read *Q*, *Q* is marked as contamination by PlsChain if |𝑄| > 3𝐿.

#### Implementation details

PlsChain is implemented in C and designated for Unix-like systems. It employs an external C library klib to construct hash tables and parse input sequencing reads. All *k*-mers are stored as 64-bit unsigned integers. Thus, the value of *k* is upper bound by 32. In addition, A light-weighted Python script is provided to post-process the per-read classification result by performing fuzzy matching against partially classified reads and grouping the reads from the same class for downstream analysis such as library diversity.

### Benchmarking PlsChain against Minimap2

#### Evaluation metrics

Precision (or positive predictive value) is defined as the total number of correct classifications, divided by the total number of attempted classification, which is also referred to classification accuracy (**Figure S4**). Recall (or sensitivity) is defined as the total number of correct classifications, divided by the total number of reads to be processed. F1-Score is defined as the harmonic mean of precision and recall (**Figure S5**).

#### Performance with real datasets

To evaluate the classification accuracy of PlsChain on real datasets, we individually prepared and assembled 3 multiplexed samples (mixture 1, 2, and 3) from 7 unique sequences with varying composition ratio. PlsChain (commit ID: 0fc9da6) was benchmarked against the minimap2 (2.28-r1209) alignment method, where all reads are aligned onto all 6348 combinations and classified based on their primary aligned combinations. In addition, for each combination, we duplicated and concatenated the sequence adjacently before alignment to account for the circular genome structure of plasmid. For the indexing step, we use k=15 (by default) for PlsChain and default settings for Minimap2 with the “secondary=no” and “-t 16” options. A read is considered to be classified by Minimap2 if it is uniquely aligned to a single reference. We refer to the fuzzy match mode for PlsChain as PlsChain-Fuzzy.

**Figure S4** quantifies the classification accuracy against read length for PlsChain, PlsChain-Fuzzy and Minimap2. It is well noted that PlsChain is more stringent on classifying reads with insufficient read length. Minimap2 tends to classify more reads than PlsChain but suffers from low classification accuracy with high false-positive classification rate. Although It is hard to observe the performance difference between PlsChain and PlsChain-Fuzzy on the plot (**Figure S4**), For mixture 1, 2, and 3 datasets, PlsChain-Fuzzy turns 317, 272, and 358 partial-classifications into fully-classification. It introduces 291, 256, and 329 more true-positive classifications while introducing 26, 16, and 29 more false-positive classifications.

#### Performance with varying k-mer size on simulated datasets

To investigate the performance with varying *k*-mer size (a parameter common to both PlsChain and Minimap2), we simulated 2000 reads against 5 randomly selected combinations out of an ɑMF1 library of expression cassettes (6348 combinations). The number of simulated reads per combination is 400. 5% sequencing errors (1.25% insertion rate, 1.25% deletion rate, 1.25% mismatch rate, and 1.25% N’s rate) were introduced per read. Cyclic shift operations are performed on simulated reads in 1*/*2 probability to emulate the circular genome structure of plasmid. For each simulated read *Q*, the probability to select a nucleotide position *p* ∈ [0,|*Q*|) is 1*/*|*Q*|, and a cyclic shift operation is performed by shifting the first *q* nucleotides on *Q* to the end of *Q*. we conducted the experiments with increasing *k*=15 up to 32 (minimap2 is upper bounded by *k* = 28) and 10 trials per *k* with different random seed to avoid biases towards particular random seed.

Figure S5 summarises the precision, recall, and F1 score of PlsChain and Minimap2 on simulated datasets with varying *k-*mer size. PlsChain (PlsChain-Fuzzy) can maintain the precision rate above 99.797% (99.799%) crossing various *k-*mer sizes. When *k*=15, both PlsChain and PlsChain-Fuzzy can achieve the recall rate as 99.55% and F1 score as 99.6% and 99.75%. When *k* is large, there is decreased probability of finding informative *k*-mers which leads to increased partial-classification rate. Minimap2 has good precision (96.75%) and recall (96.75%) when *k*=15. However, Minimap2 scales poorly with increasing *k*-mer size on precision as shown on the plot. The result reflects that traditional alignment methods may not work for classifying whole genome sequences with good classification accuracy. It’s well noted that PlsChain-Fuzzy achieves much better recall and F1-score compared to PlsChain with increasing *k*-mer size while maintaining extremely high precision rate. Hence, PlsChain-Fuzzy may be scalable to longer plasmid sequences with increasing *k*-mer size.

#### Software and Resource Usage

All the experiments were run under the National Computational Infrastructure (NCI) Gadi supercomputer by submitting jobs to the Gadi biodev queue with the default job dependencies. The allocated RAM size was limited to 500 GB and Wall time was limited to 50 h. CPU time and peak memory (maximum resident set size) is monitored by GNU time command.

Table S6 summarises the CPU time, Wall time, and peak memory on the Mixture 1-3 datasets. Thanks to the unique k-mer matching and hash table, PlsChain can process the mixture datasets (∼0.1GB in gzipped format) in roughly 6∼8 seconds with peak memory roughly 0.2GB, whereas Minimap2 takes roughly 3 hours with peak memory roughly 11GB. Since PlsChain-Fuzzy only takes an additional time overhead of roughly 0.5 seconds compared to PlsChain, we didn’t account for the actual performance of PlsChain-Fuzzy in the table. In the actual implementation, users can obtain outputs from both PlsChain and PlsChain-Fuzzy concurrently after executing the PlsChain post-processing script after performing PlsChain classification.

## Supplemental information

Supplementary File 1 - Supplementary Figures, tables and associated descriptions.

Supplementary File 2 - high resolution heat maps of PlsChain classification of ɑMF1 and ɑMF3 Nanopore library sequencing.

Supplementary file 3 - raw signal peptide sequences used for analysis and the 46 most diverse signal peptides selected for use in combinatorial Golden Gate assembly (also available within a public Benchling folder). File also includes MEGA, SVD and tNSE analysis python code and description file

Supplementary file 4 - sequences of all primers, novel C-terminal tags and raw ELISA vs FACS data (used to create Figures 3C and 4a, subset of data is shown in Table 1).

[Public Benchling project](https://benchling.com/x_josbr/f_/wrOmoFXE-protein_secretion_biosensor_ms_2024/), containing plasmid maps, assemblies and sequences.

## References

1. 1. Liu, L., Wang, J., Rosenberg, D., Zhao, H., Lengyel, G., and Nadel, D. (2018). Fermented beverage and food storage in 13,000 y-old stone mortars at Raqefet Cave, Israel: Investigating Natufian ritual feasting. J. Archaeol. Sci. Rep. 21, 783–793. 10.1016/j.jasrep.2018.08.008.

2. Wang, G., Huang, M., and Nielsen, J. (2017). Exploring the potential of Saccharomyces cerevisiae for biopharmaceutical protein production. Curr. Opin. Biotechnol. 48, 77–84. 10.1016/j.copbio.2017.03.017.

3. Huang, M., Bao, J., Hallström, B.M., Petranovic, D., and Nielsen, J. (2017). Efficient protein production by yeast requires global tuning of metabolism. Nat. Commun. 8. 10.1038/s41467-017-00999-2.

4. Püllmann, P., and Weissenborn, M.J. (2021). Improving the Heterologous Production of Fungal Peroxygenases through an Episomal Pichia pastoris Promoter and Signal Peptide Shuffling System. ACS Synth. Biol. 10, 1360–1372. 10.1021/acssynbio.0c00641.

5. Kulagina, N., Besseau, S., Godon, C., Goldman, G.H., Papon, N., and Courdavault, V. (2021). Yeasts as Biopharmaceutical Production Platforms. Front. Fungal Biol. 2, 733492. 10.3389/ffunb.2021.733492.

6. Dupuis, J.H., Cheung, L.K.Y., Newman, L., Dee, D.R., and Yada, R.Y. (2023). Precision cellular agriculture: The future role of recombinantly expressed protein as food. Compr. Rev. Food Sci. Food Saf. 22, 882–912. 10.1111/1541-4337.13094.

7. Barone, G.D., Emmerstorfer-Augustin, A., Biundo, A., Pisano, I., Coccetti, P., Mapelli, V., and Camattari, A. (2023). Industrial Production of Proteins with Pichia pastoris—Komagataella phaffii. Biomolecules 13, 441. 10.3390/biom13030441.

8. Püllmann, P., Knorrscheidt, A., Münch, J., Palme, P.R., Hoehenwarter, W., Marillonnet, S., Alcalde, M., Westermann, B., and Weissenborn, M.J. (2021). A modular two yeast species secretion system for the production and preparative application of unspecific peroxygenases. Commun. Biol. 4. 10.1038/s42003-021-02076-3.

9. Bae, J.H., Sung, B.H., Kim, H.J., Park, S.H., Lim, K.M., Kim, M.J., Lee, C.R., and Sohn, J.H. (2015). An Efficient Genome-Wide Fusion Partner Screening System for Secretion of Recombinant Proteins in Yeast. Sci. Rep. 5, 1–15. 10.1038/srep12229.

10. Delic, M., Valli, M., Graf, A.B., Pfeffer, M., Mattanovich, D., and Gasser, B. (2013). The secretory pathway: Exploring yeast diversity. FEMS Microbiol. Rev. 37, 872–914. 10.1111/1574-6976.12020.

11. Owji, H., Nezafat, N., Negahdaripour, M., Hajiebrahimi, A., and Ghasemi, Y. (2018). A comprehensive review of signal peptides: Structure, roles, and applications. Eur. J. Cell Biol. 97, 422–441. 10.1016/j.ejcb.2018.06.003.

12. Nyathi, Y., Wilkinson, B.M., and Pool, M.R. (2013). Co-translational targeting and translocation of proteins to the endoplasmic reticulum. Biochim. Biophys. Acta 1833, 2392–2402. 10.1016/j.bbamcr.2013.02.021.

13. Chaudhuri, B., Steube, K., and Stephan, C. (1992). The pro-region of the yeast prepro-α-factor is essential for membrane translocation of human insulin-like growth factor 1 *in vivo*. Eur. J. Biochem. 206, 793–800. 10.1111/j.1432-1033.1992.tb16986.x.

14. Fitzgerald, I., and Glick, B.S. (2014). Secretion of a foreign protein from budding yeasts is enhanced by cotranslational translocation and by suppression of vacuolar targeting. Microb. Cell Factories 13, 1–12. 10.1186/s12934-014-0125-0.

15. Rakestraw, J.A., Sazinsky, S.L., Piatesi, A., Antipov, E., and Wittrup, K.D. (2009). Directed evolution of a secretory leader for the improved expression of heterologous proteins and full-length antibodies in Saccharomyces cerevisiae. Biotechnol. Bioeng. 103, 1192–1201. 10.1002/bit.22338.

16. von Heijne, G. (1984). How signal sequences maintain cleavage specificity. J. Mol. Biol. 173, 243–251. 10.1016/0022-2836(84)90192-X.

17. Choo, K.H., and Ranganathan, S. (2008). Flanking signal and mature peptide residues influence signal peptide cleavage. BMC Bioinformatics 9, S15. 10.1186/1471-2105-9-S12-S15.

18. Snapp, E.L., McCaul, N., Quandte, M., Cabartova, Z., Bontjer, I., Källgren, C., Nilsson, I., Land, A., Von Heijne, G., Sanders, R.W., et al. (2017). Structure and topology around the cleavage site regulate post-translational cleavage of the HIV-1 gp160 signal peptide. eLife 6, 1–25. 10.7554/eLife.26067.

19. Williams, E.J.B., Pal, C., and Hurst, L.D. (2000). The molecular evolution of signal peptides. Gene 253, 313–322. 10.1016/S0378-1119(00)00233-X.

20. Tirincsi, A., Sicking, M., Hadzibeganovic, D., Haßdenteufel, S., and Lang, S. (2022). The molecular biodiversity of protein targeting and protein transport related to the endoplasmic reticulum. Int. J. Mol. Sci. 23. 10.3390/ijms23010143.

21. Obst, U., Lu, T.K., and Sieber, V. (2017). A Modular Toolkit for Generating Pichia pastoris Secretion Libraries. ACS Synth. Biol. 6, 1016–1025. 10.1021/acssynbio.6b00337.

22. Liu, Z., Tyo, K.E.J., Martínez, J.L., Petranovic, D., and Nielsen, J. (2012). Different expression systems for production of recombinant proteins in *Saccharomyces cerevisiae*. Biotechnol. Bioeng. 109, 1259–1268. 10.1002/bit.24409.

23. Sosa-Carrillo, S., Galez, H., Napolitano, S., Bertaux, F., and Batt, G. (2023). Maximizing protein production by keeping cells at optimal secretory stress levels using real-time control approaches. Nat. Commun. 14, 3028. 10.1038/s41467-023-38807-9.

24. Peng, K., Kroukamp, H., Pretorius, I.S., and Paulsen, I.T. (2021). Yeast Synthetic Minimal Biosensors for Evaluating Protein Production. ACS Synth. Biol. 10, 1640–1650. 10.1021/acssynbio.0c00633.

25. Thak, E.J., Yoo, S.J., Moon, H.Y., and Kang, H.A. (2020). Yeast synthetic biology for designed cell factories producing secretory recombinant proteins. FEMS Yeast Res. 20, 1–17. 10.1093/femsyr/foaa009.

26. Mori, A., Hara, S., Sugahara, T., Kojima, T., Iwasaki, Y., Kawarasaki, Y., Sahara, T., Ohgiya, S., and Nakano, H. (2015). Signal peptide optimization tool for the secretion of recombinant protein from Saccharomyces cerevisiae. J. Biosci. Bioeng. 120, 518–525. 10.1016/j.jbiosc.2015.03.003.

27. Peng, C., Guo, Y., Ren, S., Li, C., Liu, F., and Lu, F. (2022). SPSED: A Signal Peptide Secretion Efficiency Database. Front. Bioeng. Biotechnol. 9, 1–4. 10.3389/fbioe.2021.819789.

28. Grasso, S., Dabene, V., Hendriks, M.M.W.B., Zwartjens, P., Pellaux, R., Held, M., Panke, S., Van Dijl, J.M., Meyer, A., and Van Rij, T. (2023). Signal Peptide Efficiency: From High-Throughput Data to Prediction and Explanation. ACS Synth. Biol. 12, 390–404. 10.1021/acssynbio.2c00328.

29. Huang, M., Wang, G., Qin, J., Petranovic, D., and Nielsen, J. (2018). Engineering the protein secretory pathway of Saccharomyces cerevisiae enables improved protein production. Proc. Natl. Acad. Sci. U. S. A. 115, E11025–E11032. 10.1073/pnas.1809921115.

30. Johansson, S.A., Dulermo, T., Jann, C., Smith, J.D., Pryszlak, A., Pignede, G., Schraivogel, D., Colavizza, D., Desfougères, T., Rave, C., et al. (2023). Large scale microfluidic CRISPR screening for increased amylase secretion in yeast. Lab. Chip 23, 3704–3715. 10.1039/D3LC00111C.

31. Tashiro, K., Nakano, T., and Honjo, T. (1997). Signal sequence trap. Expression cloning method for secreted proteins and type 1 membrane proteins. Methods Mol. Biol. Clifton NJ 69, 203–219. 10.1385/0-89603-383-x:203.

32. Shaw, W.M., Yamauchi, H., Mead, J., Gowers, G.O.F., Bell, D.J., Öling, D., Larsson, N., Wigglesworth, M., Ladds, G., and Ellis, T. (2019). Engineering a Model Cell for Rational Tuning of GPCR Signaling. Cell 177, 782–796.e27. 10.1016/j.cell.2019.02.023.

33. Michaelis, S., and Barrowman, J. (2012). Biogenesis of the Saccharomyces cerevisiae Pheromone a-Factor, from Yeast Mating to Human Disease. Microbiol. Mol. Biol. Rev. 76, 626–651. 10.1128/mmbr.00010-12.

34. Sleep, D., Belfield, G.P., and Goodey, A.R. (1990). The Secretion of Human Serum Albumin from the Yeast Saccharomyces cerevisiae Using Five Different Leader Sequences. Bio/Technology 8, 42–46. 10.1038/nbt0190-42.

35. Kang, H.A., Choi, E.-S., Hong, W.-K., Kim, J.-Y., Ko, S.-M., Sohn, J.-H., and Rhee, S.K. (2000). Proteolytic stability of recombinant human serum albumin secreted in the yeast Saccharomyces cerevisiae. Appl. Microbiol. Biotechnol. 53, 575–582. 10.1007/s002530051659.

36. Lee, M.E., DeLoache, W.C., Cervantes, B., and Dueber, J.E. (2015). A Highly Characterized Yeast Toolkit for Modular, Multipart Assembly. ACS Synth. Biol. 4, 975–986. 10.1021/sb500366v.

37. Nett, J.H., Stadheim, T.A., Li, H., Bobrowicz, P., Hamilton, S.R., Davidson, R.C., Choi, B., Mitchell, T., Bobrowicz, B., Rittenhour, A., et al. (2011). A combinatorial genetic library approach to target heterologous glycosylation enzymes to the endoplasmic reticulum or the Golgi apparatus of *Pichia pastoris*. Yeast 28, 237–252. 10.1002/yea.1835.

38. Otto, M., Skrekas, C., Gossing, M., Gustafsson, J., Siewers, V., and David, F. (2021). Expansion of the Yeast Modular Cloning Toolkit for CRISPR-Based Applications, Genomic Integrations and Combinatorial Libraries. ACS Synth. Biol. 10, 3461–3474. 10.1021/acssynbio.1c00408.

39. Naseri, G., and Koffas, M.A.G. (2020). Application of combinatorial optimization strategies in synthetic biology. Nat. Commun. 11, 2446. 10.1038/s41467-020-16175-y.

40. Heiman, M.G., Engel, A., and Walter, P. (2007). The Golgi-resident protease Kex2 acts in conjunction with Prm1 to facilitate cell fusion during yeast mating. J. Cell Biol. 176, 209–222. 10.1083/jcb.200609182.

41. Almagro Armenteros, J.J., Tsirigos, K.D., Sønderby, C.K., Petersen, T.N., Winther, O., Brunak, S., von Heijne, G., and Nielsen, H. (2019). SignalP 5.0 improves signal peptide predictions using deep neural networks. Nat. Biotechnol. 37, 420–423. 10.1038/s41587-019-0036-z.

42. Katoh, K., and Standley, D.M. (2013). MAFFT Multiple Sequence Alignment Software Version 7: Improvements in Performance and Usability. Mol. Biol. Evol. 30, 772–780. 10.1093/molbev/mst010.

43. Tamura, K., Stecher, G., and Kumar, S. (2021). MEGA11: Molecular Evolutionary Genetics Analysis Version 11. Mol. Biol. Evol. 38, 3022–3027. 10.1093/molbev/msab120.

44. Shaner, M.C., Blair, I.M., and Schneider, T.D. (1993). Sequence logos: a powerful, yet simple, tool. In [1993] Proceedings of the Twenty-sixth Hawaii International Conference on System Sciences, pp. 813–821 vol.1. 10.1109/HICSS.1993.270609.

45. Okabayashi, K., Nakagawa, Y., Hayasuke, N., Ohi, H., Miura, M., Ishida, Y., Shimizu, M., Murakami, K., Hirabayashi, K., Minamino, H., et al. (1991). Secretory Expression of the Human Serum Albumin Gene in the Yeast, Saccharomyces cerevisiae. J. Biochem. (Tokyo) 110, 103–110. 10.1093/oxfordjournals.jbchem.a123527.

46. Mallem, M., Warburton, S., Li, F., Shandil, I., Nylen, A., Kim, S., Jiang, Y., Meehl, M., d’Anjou, M., Stadheim, T.A., et al. (2014). Maximizing recombinant human serum albumin production in a Mut ^s^ *Pichia pastoris* strain. Biotechnol. Prog. 30, 1488–1496. 10.1002/btpr.1990.

47. Sikkema, A.P., Tabatabaei, S.K., Lee, Y.-J., Lund, S., and Lohman, G.J.S. (2023). High-Complexity One-Pot Golden Gate Assembly. Curr. Protoc. 3, e882. 10.1002/cpz1.882.

48. Edmonds, J. (1967). Optimum branchings. J. Res. Natl. Bur. Stand. Sect. B Math. Math. Phys. 71B, 233. 10.6028/jres.071B.032.

49. Li, H. (2018). Minimap2: pairwise alignment for nucleotide sequences. Bioinformatics 34, 3094–3100. 10.1093/bioinformatics/bty191.

50. Gronchi, N., De Bernardini, N., Cripwell, R.A., Treu, L., Campanaro, S., Basaglia, M., Foulquié-Moreno, M.R., Thevelein, J.M., Van Zyl, W.H., Favaro, L., et al. (2022). Natural Saccharomyces cerevisiae Strain Reveals Peculiar Genomic Traits for Starch-to-Bioethanol Production: the Design of an Amylolytic Consolidated Bioprocessing Yeast. Front. Microbiol. 12, 768562. 10.3389/fmicb.2021.768562.

51. Ito, Y., Ishigami, M., Terai, G., Nakamura, Y., Hashiba, N., Nishi, T., Nakazawa, H., Hasunuma, T., Asai, K., Umetsu, M., et al. (2022). A streamlined strain engineering workflow with genome-wide screening detects enhanced protein secretion in Komagataella phaffii. Commun. Biol. 5, 561. 10.1038/s42003-022-03475-w.

52. Li, F., Chen, Y., Qi, Q., Wang, Y., Yuan, L., Huang, M., Elsemman, I.E., Feizi, A., Kerkhoven, E.J., and Nielsen, J. (2022). Improving recombinant protein production by yeast through genome-scale modeling using proteome constraints. Nat. Commun. 13, 2969. 10.1038/s41467-022-30689-7.

53. Gietz, R.D., and Schiestl, R.H. (2007). Microtiter plate transformation using the LiAc/SS carrier DNA/PEG method. Nat. Protoc. 2, 5–8. 10.1038/nprot.2007.16.

54. Novák, V. (1984). Cuts in cyclically ordered sets. Czechoslov. Math. J. 34, 322–333.

55. Bzikadze, A.V., and Pevzner, P.A. (2020). Automated assembly of centromeres from ultra-long error-prone reads. Nat. Biotechnol. 38, 1309–1316. 10.1038/s41587-020-0582-4.

