## Supplementary file 1 for "High-throughput optimisation of protein secretion in yeast via an engineered biosensor"

**Supplementary Data File 1**

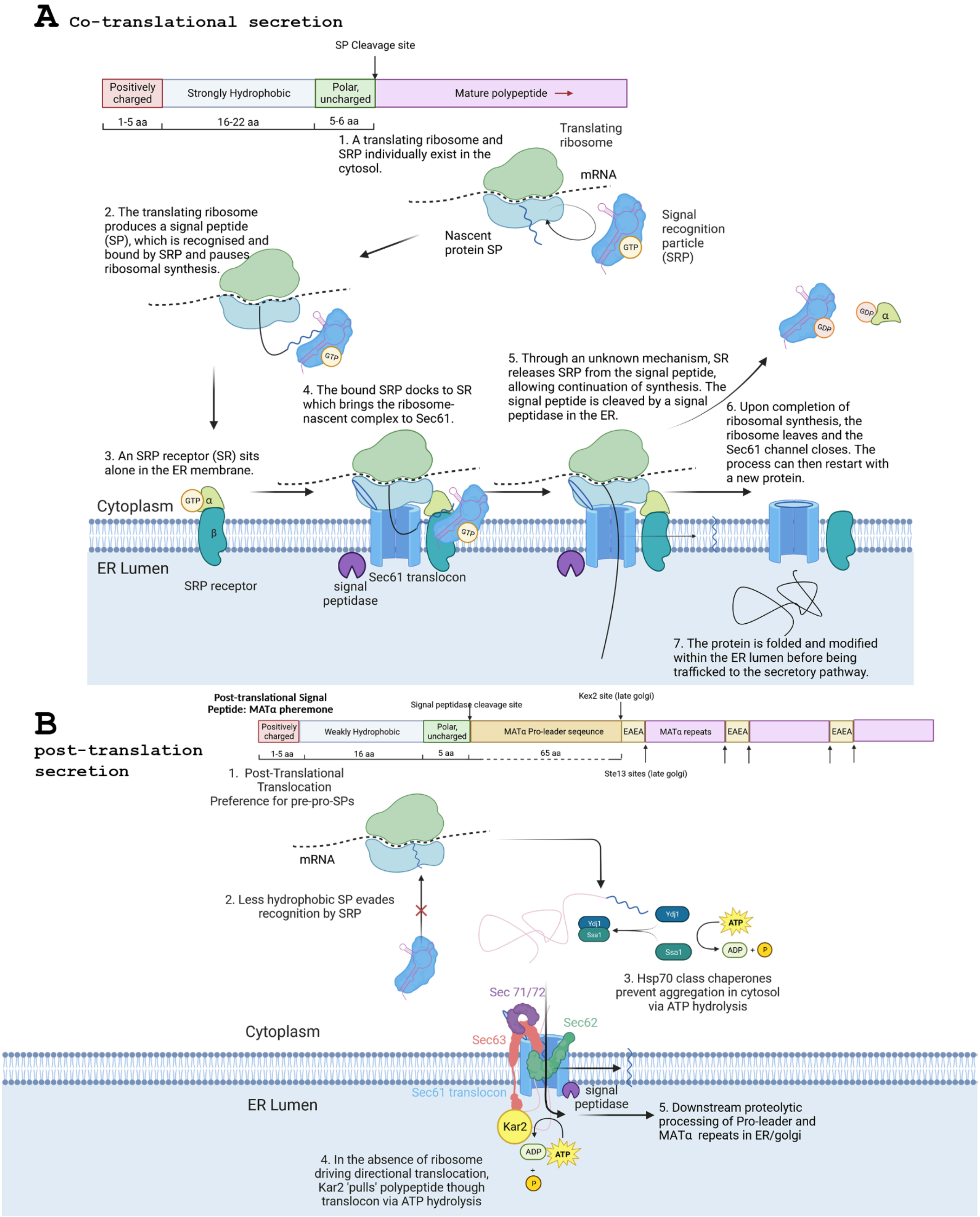


**Figure S1 Schematic comparison of co- and post-translational secretion mechanisms in yeast.**

A) Co-translational signal peptides consists of a N-terminal positively charged region, a hydrophobic region of 16-22 aa (corresponding to a transmembrane

$\alpha$-helix), and region of polar residues adjacent to the signal peptidase cleavage site. During translation by the ribosome, the strongly hydrophobic region of the nascent amino acid signal peptide is bound by a riboprotein complex, called the signal recognition particle (SRP). This interaction physically blocks interaction with elongation factor 2, preventing tRNA translocation, stalling translation . This allows a high affinity interaction with the SRP receptor heterodimer, which contains a integral membrane [illegible]-subunit and a membrane associated $\alpha$-subunit. The SRP receptor enables docking of the ribosome to the Sec61 translocon (a hexameric complex of subunits including Sec61), at which point translation resumes, driving translocation of the nascent polypeptide chain through the Sec61 transmembrane pore. The negatively charged lipids of the ER membrane outer leaflet strongly interact with the N-terminal positive charges of the signal peptide, resulting in a "hairpin" insertion into the translocon. The signal peptide is then cleaved by the associated signal peptidase, and a lateral gate allows partitioning of the hydrophobic sequence into the lipid bilayer where it is later marked for degradation. The remainder of the protein is then secreted into the ER lumen before being trafficked out of the cell via the golgi [10,12].

B) Post-translational secretion is analogous in many respects to cotranslational secretion with some important points of distinction. These are principally the absence of SRP which does not recognise the less hydrophobic character of post-translational signal peptides. There is also the requirement for Hsp70 class chaperones for folding and solubility prior to delivery to the Sec61 translocon which now requires additional ancillary subunits to form a heptameric complex. It is also thought that ATPase activity of the ER lumenal chaperon Kar2 provides force to 'pull' the nascent chain into the ER.[10] Figures made using Biorender.com.

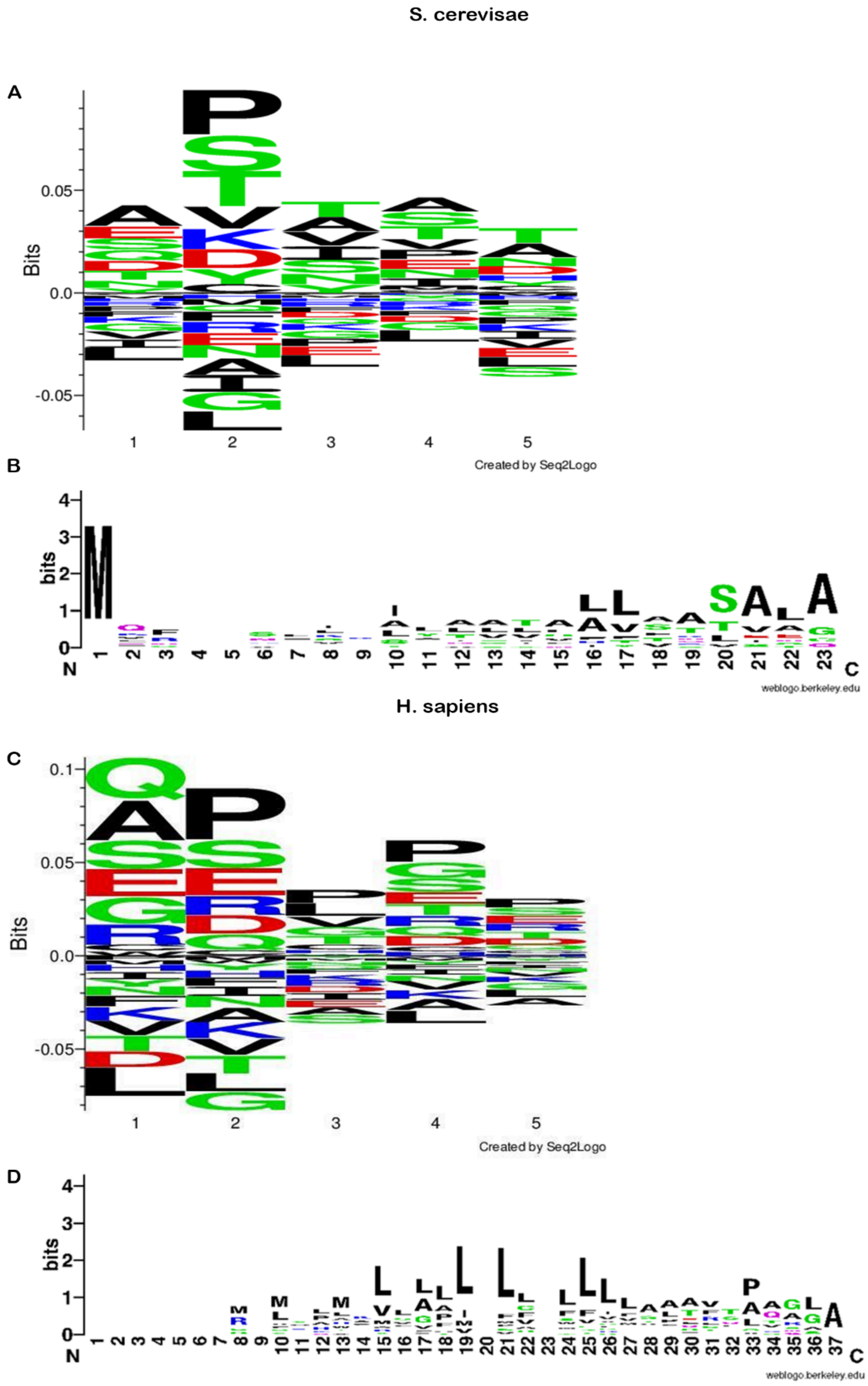
S. cerevisae
A
Bits
Created by Seq2Logo
B
bits
N
C
weblogo.berkeley.edu
H. sapiens
C
Bits
Created by Seq2Logo
D
bits
N
C
weblogo.berkeley.edu

**Figure S2**

A) SequenceLogo[44] analysis of all yeast signal peptide +5 residues downstream of the signal peptidase cleavage site predicted by Signal P,[41] showing a preference for Alanine and Proline at the +1 and +2 positions, respectively.

B) SequenceLogo[44] analysis of all of signal peptides from yeast chosen for the library qualititatively showing no clear conservation of sequence identity.

C) SequenceLogo[44] analysis of all human signal peptide +5 residues downstream of the signal peptidase cleavage site predicted by Signal P,[41] showing a preference for Alanine and Proline at the +1 and +2 positions, respectively.

D) SequenceLogo[44] analysis of all of signal peptides from humans chosen for the library, qualitatively showing a preference for Leucine residues within the hydrophobic region of these signal sequences.

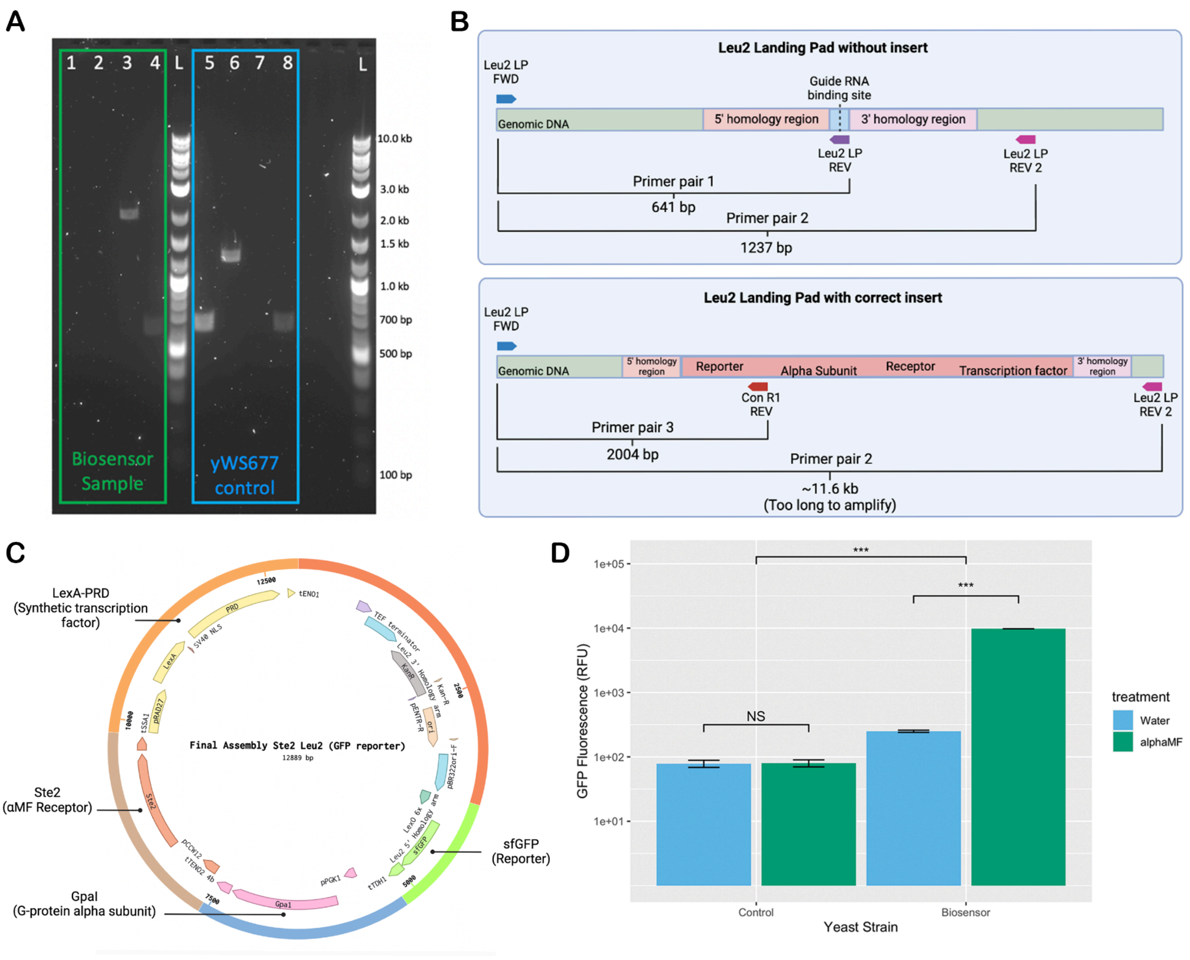


**Figure S3 PCR confirmation of yWS677 CRISPR edit to construct biosensor strain.** A) Left in Green = Biosensor sample template gDNA, Right, in Blue = yWS677 template gDNA. L = 1 kb Plus Ladder (NEB), lanes 1 and 5 were amplified using Primer Pair 1, lanes 2 and 6 were amplified using Primer Pair 2, lanes 3 and 7 were amplified using Primer Pair 3, lanes 4 and 8 were amplified using Primer Pair 4 (URA3 LP FWD + URA3 LP REV1, a primer pair known to successfully amplify

elsewhere in the yWS677 genome). Gel was run on a 1% (w/v) agarose/LAB buffer gel, visualised with 1% (w/v) SYBRSafe, at 120 V for 60 mins.
B) Diagrammatic representation of primer pairs 1, 2 and 3 binding sites and amplicons within and around the LEU2 LP region in the yWS677 genome. Primer LEU2 LP FWD binds outside the 5' homology arm of the LEU2 LP site. Primer LEU2 LP REV 1 binds directly at the guide RNA binding site/Cas9 cut site. Primer LEU2 LP REV2 binds outside the 3' homology arm of the LEU2 LP site. Primer ConR1 REV inside the donor DNA/insert sequence, just after the reporter gene. LEU2 LP FWD + LEU2 LP REV1 will amplify template gDNA that does not contain an insert and has an intact LEU2 LP guide RNA binding site (ie. The site has most likely not been cut by Cas9, or has been cut but seamlessly repaired by the cell DNA repair mechanisms). LEU2 LP FWD + LEU2 LP REV 2 will amplify template gDNA that does not contain and insert (does not require an unchanged LEU2 LP guide RNA binding site sequence (will amplify even if DNA has been cut and re-ligated, with mutations introduced at the guide RNA binding site). LEU2 LP FWD + ConR1 REV will amplify template gDNA containing the inserted biosensor genes. Figure made using BioRender.
C) Complete plasmid map of level 2 assembly used for donor DNA in CRISPR edit used to construct biosensor strain. All Golden Gate parts were from the MoClo YTK (Addgene Kit # 1000000061) or the Yeast GPCR-sensor Toolkit (Addgene Kit # 1000000157). Image made using Benchling.
D) Single cell GFP fluorescence (RFU) of control strain yWS677 compared to αMF biosensor yeast strain when treated with water or 100 mM S. cerevisiae αMF. GFP fluorescence (RFU) is represented on a log10 scale. Columns are grouped by yeast strain (Control yWS677 or αMF Biosensor) and coloured by treatment ('Water' treatment represented in blue and 'αMF' treatment represented in green). Result of two-way ANOVA represented by square bracket across yeast strains. Results of post hoc reverse pairwise comparisons between treatments within each yeast strain represented square bracket across treatments. Significance indicated by '***' = $p \le 0.001$ (highly significant), '**' = $p<0.01$ (moderately significant), '*' = $p<0.05$ (significant), 'NS' = $p>0.05$ (not significant).

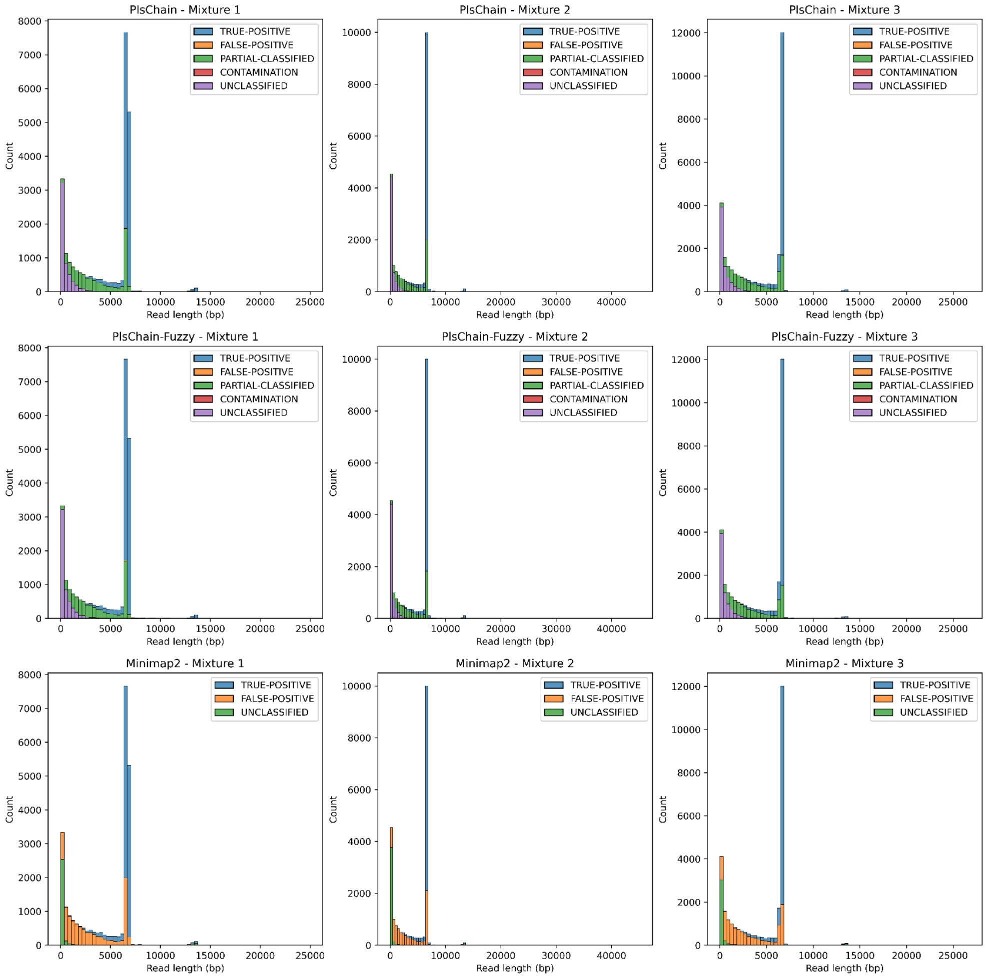


### Figure S4 Evaluation of Classification Accuracy on Real Mixture 1-3 Datasets

Histograms quantifying the classification accuracy against read length for PlsChain, PlsChain-Fuzzy and Minimap2. Different classification states (true-positive classification, false-positive classification, partial-classification, contamination, and unclassified) for PlsChain (PlsChain-Fuzzy) are labelled in different colours. Different classification states (true-positive classification, false-positive classification, and unclassified) for Minimap2 are labelled in different colours.

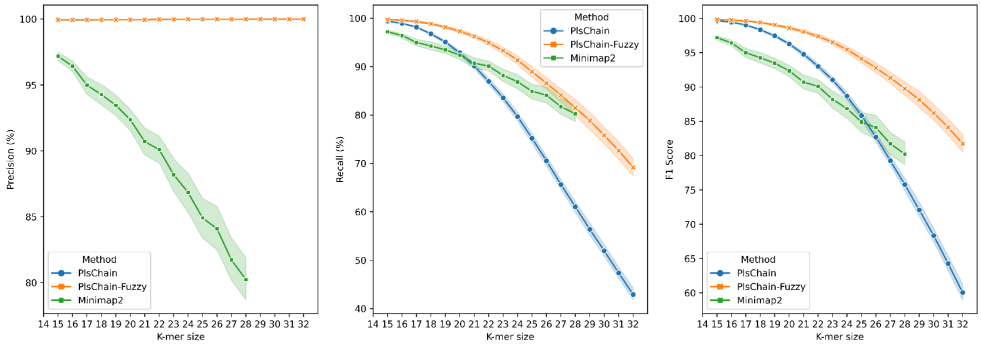


### Figure S5 Evaluation of Classification Accuracy on Simulated Datasets

Line plot quantifying the classification accuracy with increasing *k*-mer size (15 up to 32) for PlsChain, PlsChain-Fuzzy and Minimap2. Precision (left), Recall (middle), and F1-score (right) for PlsChain (blue), PlsChain-fuzzy (orange), and Minimap2 (green) are measured against simulated datasets. For every *k*-mer size, 10 simulated datasets generated by 10 different random seeds are applied for benchmarking.

| Minimap2 | Indexing | Mixture1 | Mixture2 | Mixture3 |
|---|---|---|---|---|
| User (sec) | 3.14 | 193708.84 | 144417.65 | 209213.56 |
| Sys (sec) | 0.56 | 3410.85 | 2553.29 | 3653.46 |
| Wall (sec) | 2.04 | 12355 | 9223 | 13340 |
| Peak Mem (kB) | 486448 | 11271936 | 12031952 | 11379112 |

| PlsChain | Indexing | Mixture1 | Mixture2 | Mixture3 |
|---|---|---|---|---|
| User (sec) | 0.05 | 7.64 | 5.88 | 8.1 |
| Sys (sec) | 1.03 | 0.21 | 0.17 | 0.23 |
| Wall (sec) | 1.24 | 7.88 | 6.18 | 8.43 |
| Peak Mem (kB) | 8776 | 180176 | 143860 | 192480 |

### Table S6 CPU Time and Memory Usage on Real Mixture 1-3 Datasets

CPU Time and peak memory usage of PlsChain(PlsChain-Fuzzy) and Minimap2 on Mixture 1-3 Datasets. Peak memory refers to the maximum resident set size.
