## Supplementary file 2 for "High-throughput optimisation of protein secretion in yeast via an engineered biosensor": Supplementary File 2 Description.docx

Supplementary File 2 Description – heatmap visualisation of PlsChain-Fuzzy (See method details) assembly of long read Nanopore sequencing of combinatorial golden gate assemblies. Each heatmap is accompanied by a .csv tabulation of reads corresponding to combination. Each library was sequenced 3 times, with values shown as the average of these 3 technical repeats. Details of each repeat are given in the file heatmap_info.csv

“k15_20231106_lib” prefix = sequencing of combinatorial assembly using C-terminal 4a tag corresponding to aMF1 (see Figure 3b)

“k15_20240304_bc” prefix = sequencing of combinatorial assembly using C-terminal 4a tag corresponding to aMF3 (see Figure 3b)

Due to the dense information content of each heatmap, a portion of each is shown for demonstration purposes.

Included in this supplementary file are the datasets corresponding to the two tag design used in library assemblies analysed with ELISA and flow cytometry (k15_20231106 = aMF1 tag and k15_20240304_bc = aMF4 tag). Heatmaps for all datasets are available from our Zenodo depository (<https://zenodo.org/doi/10.5281/zenodo.10980037>)
