## Supplementary figures and images for "High-throughput optimisation of protein secretion in yeast via an engineered biosensor"

### k15_20231106_lib_heatmap.fuzzy.png

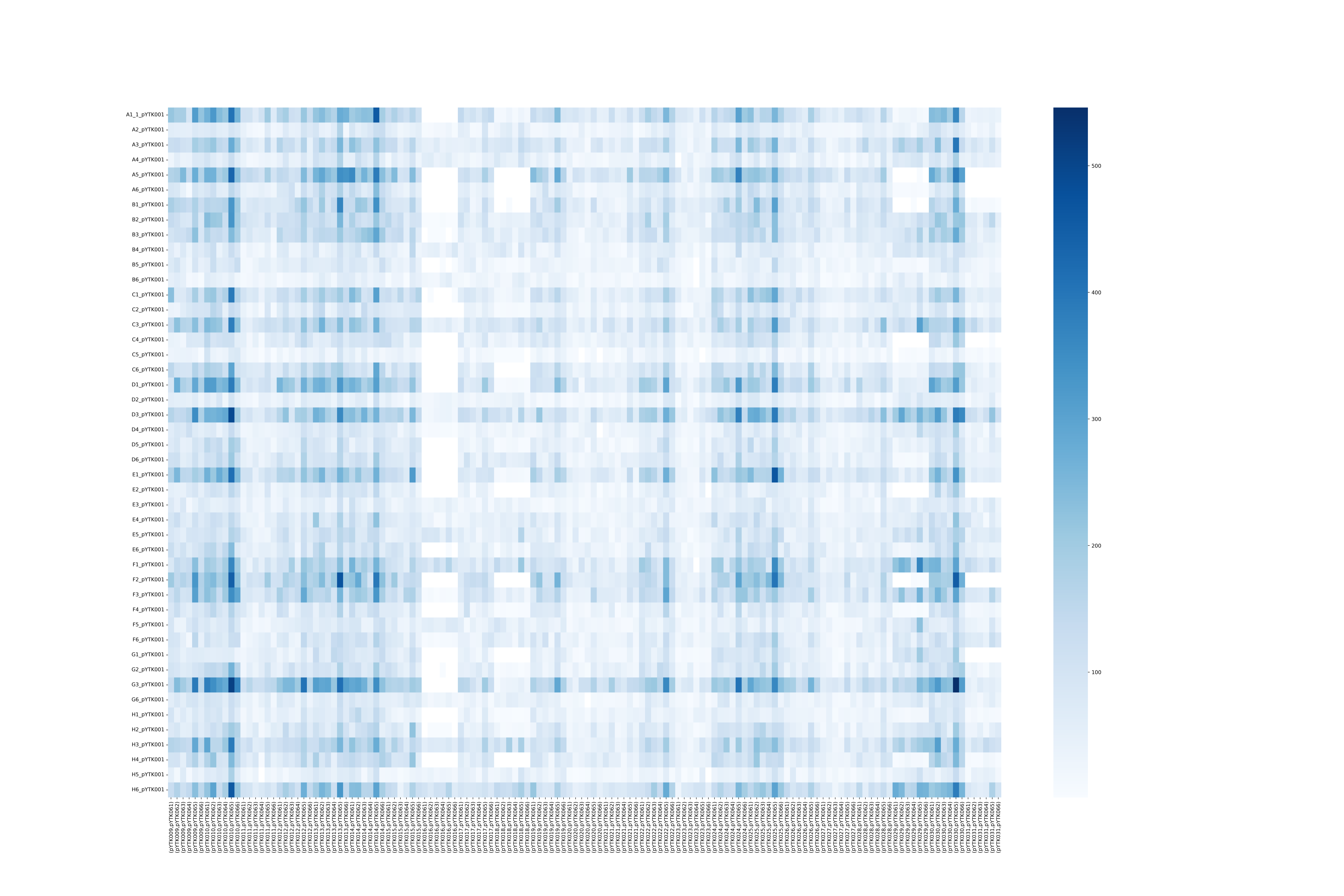

### k15_20231106_lib_heatmap_portion.fuzzy.png

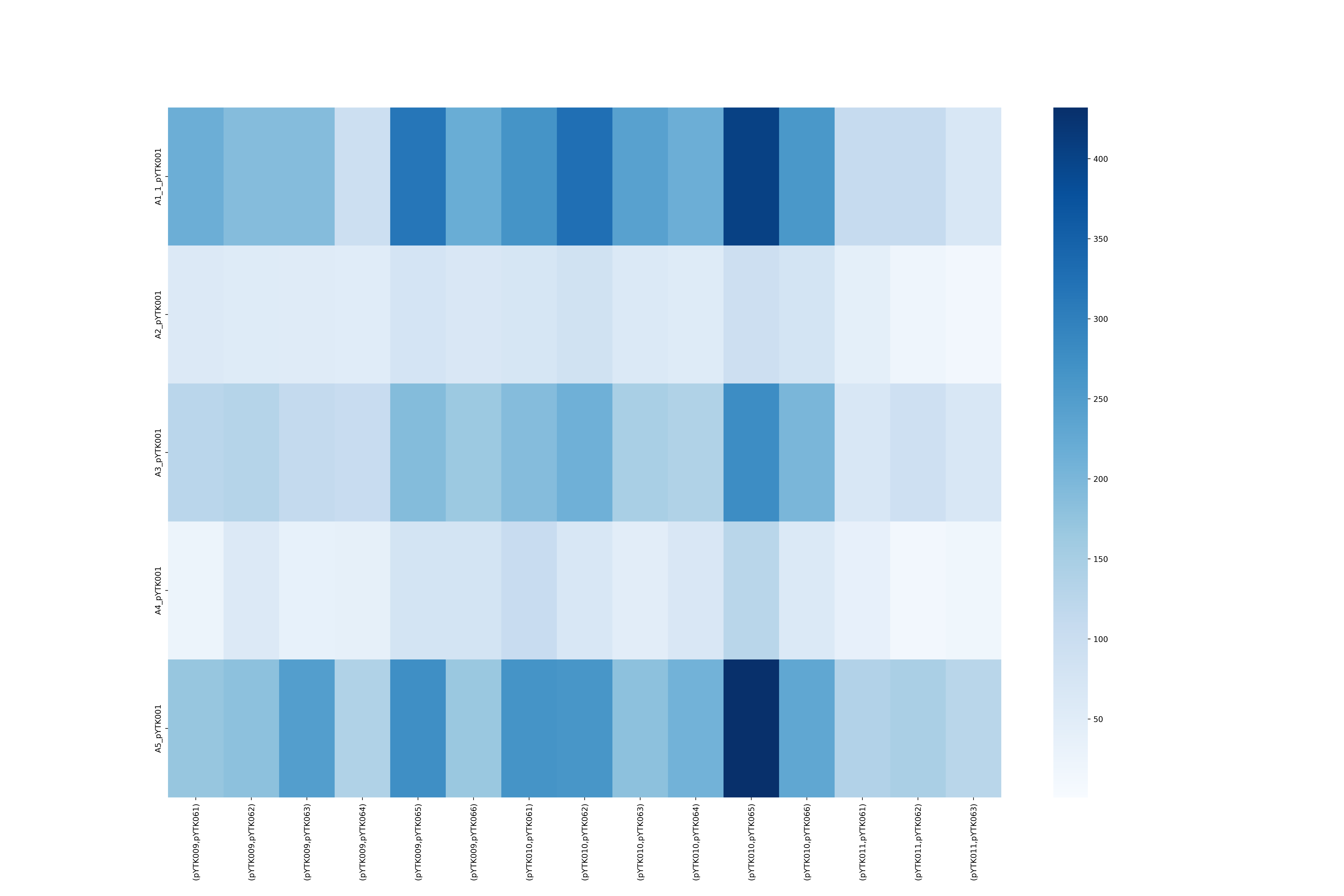

### k15_20240304_bc_heatmap.fuzzy.png

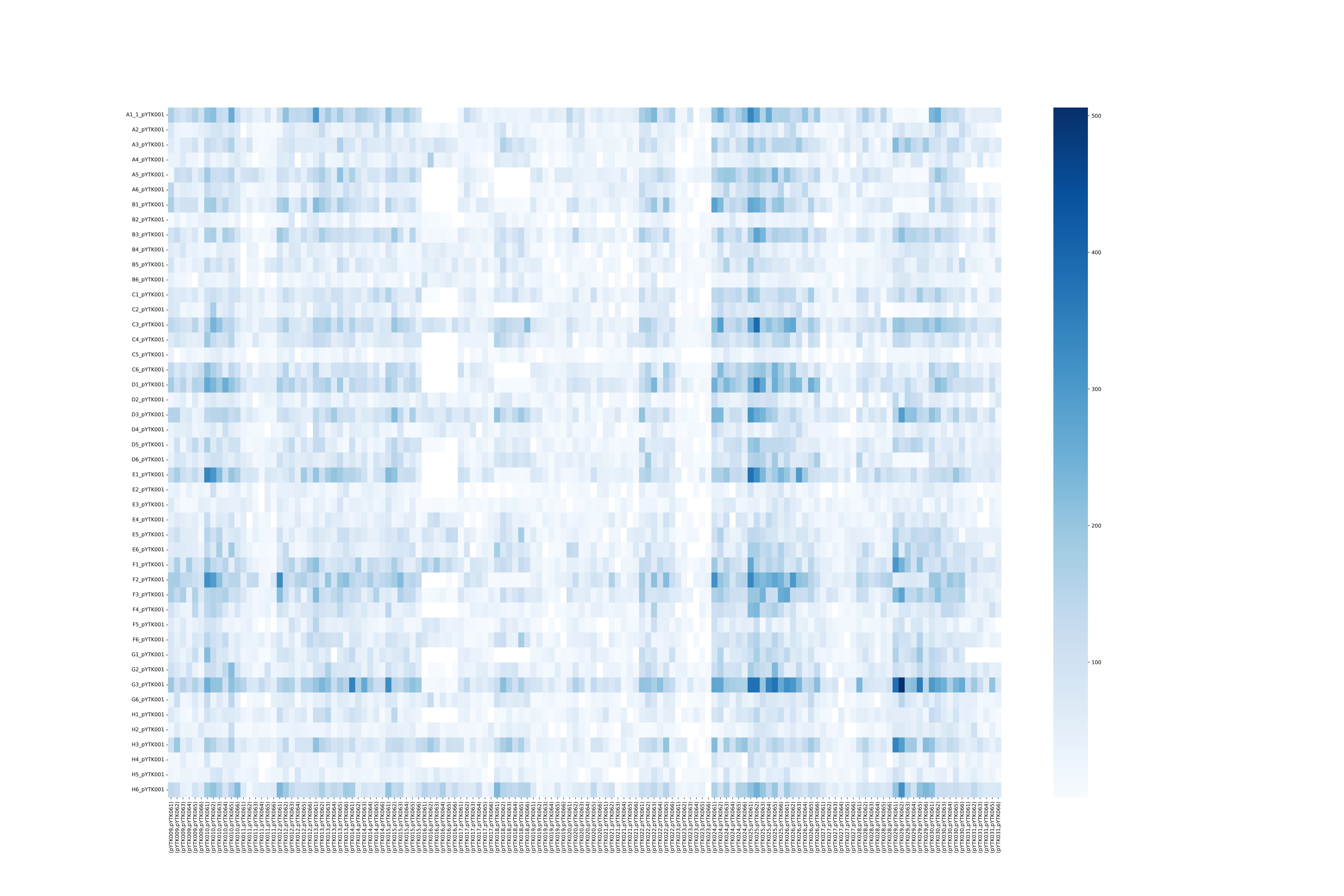

### k15_20240304_bc_heatmap_portion.fuzzy.png

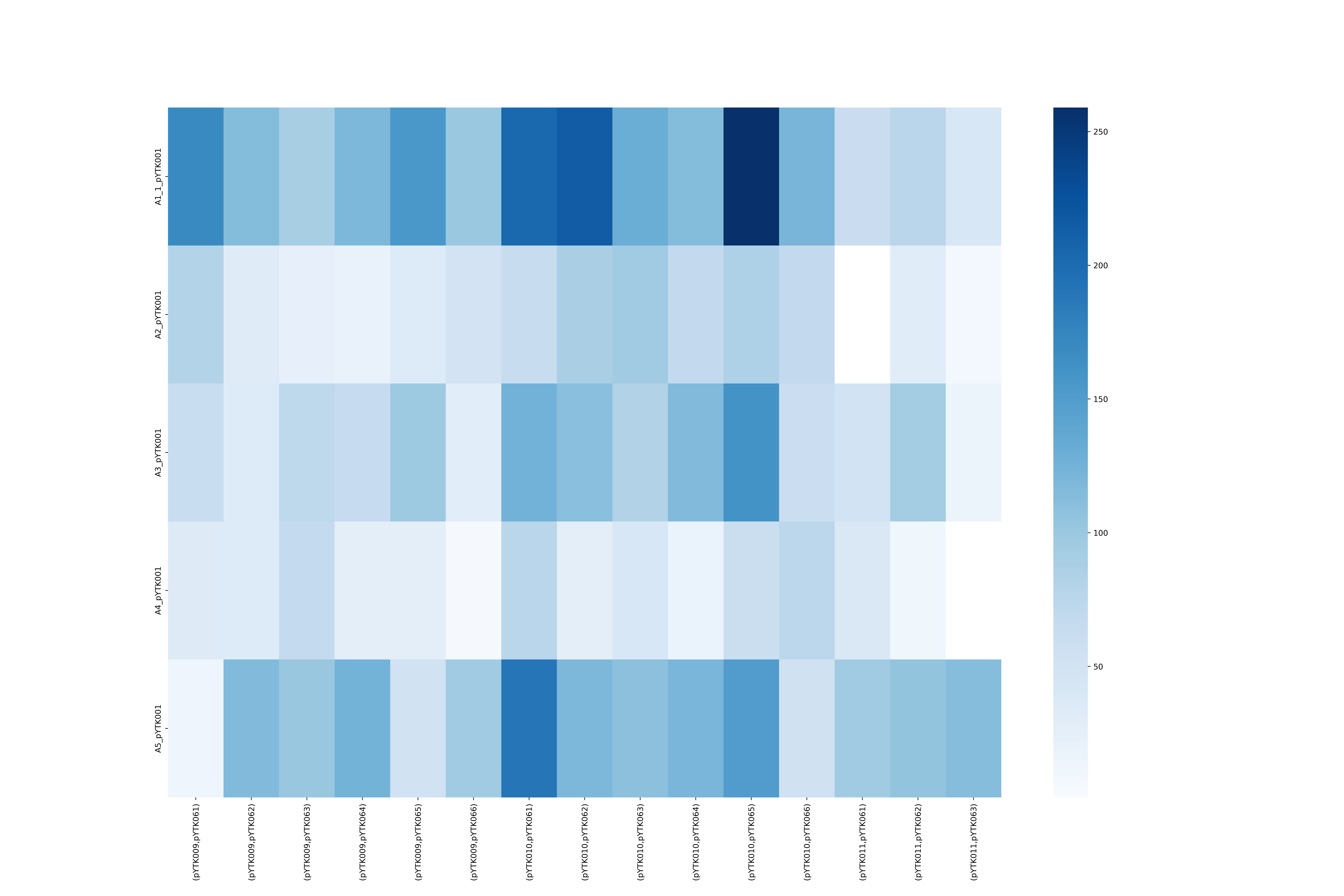
