## Supplementary file 3 for "High-throughput optimisation of protein secretion in yeast via an engineered biosensor": Intial_RFU_measurements_R code.docx

Initial 100nM aMF tests

2022-10-09

Import libraries

library(tidyverse)
library(janitor)
library(emmeans)
library(ggResidpanel)
library(ggsignif)
library(ggokabeito)

Import data

```
data<-read_csv("sorted data.csv") %>%
  clean_names() %>%
  mutate(strain = factor(strain, levels = c("Control", "Biosensor")),
         treatment = factor(treatment, levels = c("Water","alphaMF")))
glimpse(data)
## Rows: 240,000
## Columns: 3
## $ rfu       <dbl> 189.72, 572.22, 500.82, 528.36, 372.30, 444.72, 324.36, 240.…
## $ treatment <fct> Water, Water, Water, Water, Water, Water, Water, Water, Wate…
## $ strain    <fct> Biosensor, Biosensor, Biosensor, Biosensor, Biosensor, Biose…
summary(data$rfu)
##     Min.  1st Qu.   Median     Mean  3rd Qu.     Max.
## -1036.32    54.06   139.74  2546.75  1013.88 69415.10
```

Data exploration

```
ggplot(data, aes(x = strain, y = rfu, col = treatment))+
  geom_boxplot()+
  scale_y_log10() +
  geom_point(position = position_dodge(width = 0.75)) +
  theme_classic()
## Warning in self$trans$transform(x): NaNs produced
## Warning: Transformation introduced infinite values in continuous y-axis
## Warning in self$trans$transform(x): NaNs produced
## Warning: Transformation introduced infinite values in continuous y-axis
## Warning: Removed 24239 rows containing non-finite values (stat_boxplot).
## Warning: Removed 23739 rows containing missing values (geom_point).
```

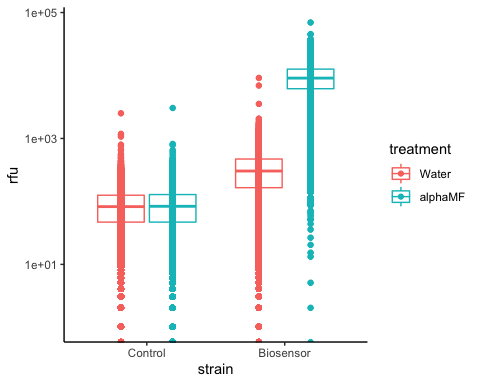

Fitting model for two-way ANOVA

```
m <- lm(rfu ~ treatment * strain, data=data)
summary(m)
##
## Call:
## lm(formula = rfu ~ treatment * strain, data = data)
##
## Residuals:
##    Min     1Q Median     3Q    Max
## -10150   -104     -9     88  59636
##
## Coefficients:
##                                  Estimate Std. Error t value Pr(>|t|)
## (Intercept)                        78.548      9.988   7.864 3.73e-15 ***
## treatmentalphaMF                    1.300     14.125   0.092    0.927
## strainBiosensor                   171.384     14.125  12.133  < 2e-16 ***
## treatmentalphaMF:strainBiosensor 9527.428     19.976 476.940  < 2e-16 ***
## ---
## Signif. codes:  0 '***' 0.001 '**' 0.01 '*' 0.05 '.' 0.1 ' ' 1
##
## Residual standard error: 2447 on 239996 degrees of freedom
## Multiple R-squared:  0.7445, Adjusted R-squared:  0.7445
## F-statistic: 2.331e+05 on 3 and 239996 DF,  p-value: < 2.2e-16
anova(m)
## Analysis of Variance Table
##
## Response: rfu
##                      Df     Sum Sq    Mean Sq F value    Pr(>F)
## treatment             1 1.3623e+12 1.3623e+12  227596 < 2.2e-16 ***
## strain                1 1.4613e+12 1.4613e+12  244134 < 2.2e-16 ***
## treatment:strain      1 1.3616e+12 1.3616e+12  227472 < 2.2e-16 ***
## Residuals        239996 1.4365e+12 5.9857e+06
## ---
## Signif. codes:  0 '***' 0.001 '**' 0.01 '*' 0.05 '.' 0.1 ' ' 1
resid_panel(m)
```

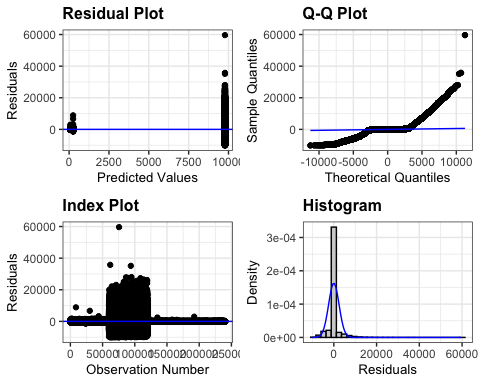

Calculating mean estimates and contrasts, with posthoc inferences.

```
emmeans(m, revpairwise ~ treatment|strain)
## $emmeans
## strain = Control:
##  treatment  emmean    SE     df lower.CL upper.CL
##  Water       78.55 9.988 239996    58.97    98.12
##  alphaMF     79.85 9.988 239996    60.27    99.42
##
## strain = Biosensor:
##  treatment  emmean    SE     df lower.CL upper.CL
##  Water      249.93 9.988 239996   230.36   269.51
##  alphaMF   9778.66 9.988 239996  9759.08  9798.24
##
## Confidence level used: 0.95
##
## $contrasts
## strain = Control:
##  contrast        estimate    SE     df t.ratio p.value
##  alphaMF - Water      1.3 14.13 239996   0.092  0.9267
##
## strain = Biosensor:
##  contrast        estimate    SE     df t.ratio p.value
##  alphaMF - Water   9528.7 14.13 239996 674.588  <.0001
```

Graphing data as estimated marginal means by treatment and strain.

```
results1 <- emmeans(m, ~ treatment|strain) %>%
  as_tibble()

finalplot <- ggplot(results1, aes(x = strain, y = emmean, fill = treatment))+
  geom_col(position = position_dodge(width = 1))+
  scale_y_log10(breaks = c(1e1, 1e2, 1e3, 1e4, 1e5),)+
  xlab("Yeast Strain")+
  ylab("GFP Fluorescence (RFU)")+

  ggtitle("GFP fluorescence of control vs. alphaMF biosensor yeast strain
          when treated with water or 100 nM alpha-MF")+
  geom_errorbar(aes(ymin=emmean - SE, ymax = emmean + SE), position = position_dodge(width = 1), width = 0.3) +

  scale_fill_okabe_ito(order = c(2,3))+

  geom_signif(comparisons=list(c("Control", "Biosensor")), annotations="***",
              y_position = 5, tip_length = 0.05, vjust=0.4) +

   geom_signif(y_position = c(2.5,4.5), xmin = c(0.75,1.75),
              xmax = c(1.25,2.25), annotation = c("NS","***"),
              tip_length = 0.05)

finalplot
```

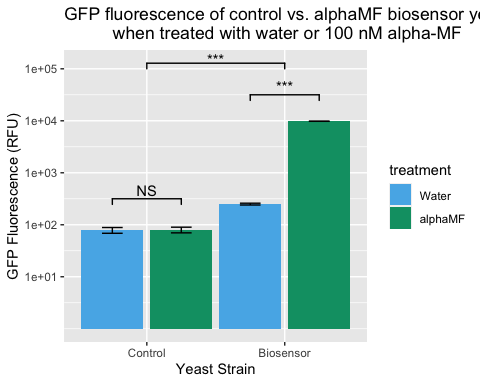

Exporting the final plot as an image.

ggsave("Final Plot.png", width = 7, height = 5)
