## Supplementary file 3 for "High-throughput optimisation of protein secretion in yeast via an engineered biosensor": mega11_to_csv.py

```
# -*- coding: utf-8 -*-
"""MEGA to CSV.ipynb

Automatically generated by Colab.

Original file is located at
    https://colab.research.google.com/drive/1dXU8pyD-2nWsJk3pmPwlLp9jb4e4HwTQ
"""

# Imports and data that we ned
import sys
import argparse
from google.colab import files

"""
Converts ugly MEGA (v11) format text files to CSV files, without restricting the number of columns.
Saves as a complete matrix with 0-distance identity diagonal
"""

def main():

    uploaded = files.upload()
    args_input = "homosapiensmega.txt"
    args_output = "output_megatocsv.csv"

    names = []
    rows = []
    seen_names = False
    seen_data = False
    with open(args_input, 'rt') as f:
        for line in f:
            if seen_names and seen_data:
                if len(line.strip()) == 0:
                    # empty line after data matrix
                    break  # we're done!
                # collect data and when done, finish up
                # separate sample names see if we're past the column headers
                name_cols = line.split(']')[0].replace('[','').strip().split()
                if len(name_cols) > 1:
                    continue  # skip the column header row
                data_cols = line.split(']')[1].strip().split()
                data_cols.append('')
                rows.append(data_cols)
                continue
            if not seen_names and not line.startswith('['):
                continue
            if not seen_names and line.startswith('['):
                seen_names = True
            if seen_names and not seen_data:
                if len(line.strip()) == 0:
                    # empty line after name section!
                    seen_data = True
                    continue
                name = line.split('#',1)[1].strip()
                names.append(name)

    with open(args_output, 'wt') as outf:
        outline = ',' + ','.join(names)
        print(outline, file=outf)
        for i, row in enumerate(rows):
            outline = names[i] + ',' + ','.join(row)
            print(outline, file=outf)

    ### done!


if __name__ == '__main__':
    main()

files.download("output_megatocsv.csv")
```
