## Supplementary file 3 for "High-throughput optimisation of protein secretion in yeast via an engineered biosensor": Supplementary file 3 description.docx

This file contains all signal peptides as .fasta files from *Eukaryota*, *Saccharomyces* and *H. sapiens* via and advanced search of the Uniprot database (performed on the 16^th^ of September, 2021):

- PTM/Processing --> signal peptide AND
- Organism --> as outlined above AND
- Subcellular localisation --> Secreted AND
- Reviewed/validated

From the 3 sequence datasets, a collection of the 23 most diverse were chose from the *Saccharomyces* and *H. sapiens* datasets respectively, following processing as described in the main text “these sets of protein sequences using SignalP 5.0 ^43^, to produce sequence sets of distinct “SP+5” sequence sets, including five residues downstream of the signal peptidase cleavage site predicted by the software. By including an additional five residues beyond the cleavage site, we aimed to conduct a weighted sequence analysis on both the SP sequences and the cleavage site itself, catering to signal peptidase processing. ^16–18^  MAFFT^44^ was used to independently align each sequence set, followed by MEGA11^45^ to generate a dataset-specific all-by-all distance matrix with expectation maximisation. Given this method provides high-dimensional embedding of the diversity in each set of SPs, we used single value decomposition (SVD) followed by t-distributed stochastic neighbour embedding (tSNE) to visualise the diversity based on relative positioning in a three-dimensional plot”

The selected 46 most diverse signal peptide sequences are saved in a subfolder in genbank format.

Please see methods sections for further details.
