## Supplementary file 3 for "High-throughput optimisation of protein secretion in yeast via an engineered biosensor": svd_and_t_sne_implemented_in_3d.py

```
# -*- coding: utf-8 -*-
"""SVD and t-SNE implemented in 3D.ipynb

Automatically generated by Colab.

Original file is located at
    https://colab.research.google.com/drive/11dtzeOKwHf9RUoKxp00YVM0N9VDgfiqm
"""

## Imports and data that we need

import numpy as np
import numpy.ma as ma
import pandas as pd
import matplotlib.pyplot as plt
import sklearn as skl
import plotly.express as px
from google.colab import files
from sklearn.decomposition import PCA, TruncatedSVD
from sklearn.manifold import TSNE
from sklearn.pipeline import make_pipeline
from scipy.sparse.linalg import svds
from functools import partial

## Input for our lower triangular distance matrix, a ".csv" file type

uploaded = files.upload()
distancematrix = pd.read_csv('homo_sapiens_distancematrix.csv', index_col=0)

print(type(distancematrix))
print(distancematrix.head)

## Convert into numpy array

dm = distancematrix
dm = dm.replace('?', np.nan)
dm = dm.replace('nan', np.nan)
dm = np.array(dm, dtype=np.double)
np.fill_diagonal(dm, np.double(0))

# print(dm.shape)

## Convert from lower triangular matrix into square matrix

inds = np.where(np.isnan(dm))
for row,col in zip(inds[0], inds[1]):
    if not np.isnan(dm[col,row]):
        dm[row,col] = dm[col,row]

# print(dm[10])

## Replaces missing vales in distance matrix with zero (expectation maximisation)

inds = np.where(np.isnan(dm))
for i, (row,col) in enumerate(zip(inds[0], inds[1])):
    dm[row,col] = np.double(0)

# print(dm[10])

## Adding indexes to the data set

dm2 = pd.DataFrame(data=dm, index=distancematrix.index.values, columns=distancematrix.columns.values)

# print(type(dm2))

## t-SNE and SVD definitions

num_TSNE=3  # needs to be less than or equal to 3 for the barnes-hut algorithm to work
metric = "precomputed"
state = 0
tsne = TSNE(n_components=num_TSNE, metric=metric, random_state=state)
svd = TruncatedSVD(n_components=12, n_iter=13, random_state=state)

## Run SVD followed by t-SNE
## adapted from https://github.com/scikit-learn/scikit-learn/issues/17916
## You can change the number values here if you want

tsne = make_pipeline(TruncatedSVD(n_components=12, n_iter=13, random_state=state), TSNE(n_iter=250, n_components=num_TSNE, random_state=state, perplexity=200))

wide_tsne = tsne.fit_transform(dm2)

## print(type(wide_tsne))

print(wide_tsne)
# print(wide_tsne.shape)

## Construct an empty dataframe with the sequence names as the index
## https://stackoverflow.com/questions/18176933/create-an-empty-data-frame-with-index-from-another-data-frame

df_index = pd.DataFrame(index = distancematrix.index)

# print(df_index)
# print(type(df_index))

## Construct a t-SNE dataframe - it will contain the 3 TSNE features for all sequences
## from https://shravan-kuchkula.github.io/nyc-airbnb-tsne-visual/#apply-t-sne

tsne_df = pd.DataFrame({'TSNE1': wide_tsne[:,0], 'TSNE2': wide_tsne[:,1],'TSNE3': wide_tsne[:,2]})
tsne_df.index = df_index.index
tsne_df.to_csv('tsne_final.csv', index=True)

files.download('tsne_final.csv')

# visualise 2d slices of 3d data

fig1 = px.scatter(tsne_df, x='TSNE1', y='TSNE2', hover_name=tsne_df.index, template='plotly', opacity=0.9, title='Columns 1 and 2')

fig1

fig2 = px.scatter(tsne_df, x='TSNE1', y='TSNE3', hover_name=tsne_df.index, template='plotly', opacity=0.9, title='Columns 1 and 3')

#fig2

fig3 = px.scatter(tsne_df, x='TSNE2', y='TSNE3', hover_name=tsne_df.index, template='plotly', opacity=0.9, title='Columns 2 and 3')

#fig3

# visualise 2d slices of 3d data

fig = plt.figure(figsize=(14,4))
ax1 = fig.add_subplot(131)
ax1.scatter(wide_tsne[:,0], wide_tsne[:,1])
ax2 = fig.add_subplot(132)
ax2.scatter(wide_tsne[:,0], wide_tsne[:,2])
ax3 = fig.add_subplot(133)
ax3.scatter(wide_tsne[:,1], wide_tsne[:,2])
plt.show()

highlighted_sequences = [
    'sp_A2RUU4_COLL1_HUMAN:0-28', 'sp_A6NGW2_STRCL_HUMAN:0-30', 'sp_O14793_GDF8_HUMAN:0-23', 'sp_O75629_CREG1_HUMAN:0-36', 'sp_P00748_FA12_HUMAN:0-24', 'sp_P13500_CCL2_HUMAN:0-28', 'sp_P24347_MMP11_HUMAN:0-38',
    'sp_P31431_SDC4_HUMAN:0-25', 'sp_P43652_AFAM_HUMAN:0-26', 'sp_P59827_BPIB4_HUMAN:0-22', 'sp_P59861_D131A_HUMAN:0-27', 'sp_P80075_CCL8_HUMAN:0-28', 'sp_P80098_CCL7_HUMAN:0-28', 'sp_Q14114_LRP8_HUMAN:0-37',
    'sp_Q16609_LPAL2_HUMAN:0-26', 'sp_Q6NUI6_CHADL_HUMAN:0-35', 'sp_Q6UW88_EPGN_HUMAN:0-25', 'sp_Q86TE4_LUZP2_HUMAN:0-24', 'sp_Q92832_NELL1_HUMAN:0-26', 'sp_Q99616_CCL13_HUMAN:0-28', 'sp_Q99645_EPYC_HUMAN:0-24',
    'sp_Q9H2A7_CXL16_HUMAN:0-34', 'sp_Q9NRA1_PDGFC_HUMAN:0-27', 'sp_Q9UKQ2_ADA28_HUMAN:0-23'
]

# Assuming you have a DataFrame 'df' with a column 'sequence' that corresponds to your t-SNE data
sequence_names = tsne_df.index.tolist()


colors = ['blue' if seq in highlighted_sequences else 'grey' for seq in sequence_names]
opacity = [1.0 if seq in highlighted_sequences else 0.2 for seq in sequence_names]

fig = plt.figure(figsize=(14,4))

ax1 = fig.add_subplot(131)
ax1.scatter(wide_tsne[:,0], wide_tsne[:,1], color=colors, alpha=opacity)
ax1.set_title('TSNE Dimensions 1 & 2')

ax2 = fig.add_subplot(132)
ax2.scatter(wide_tsne[:,0], wide_tsne[:,2], color=colors, alpha=opacity)
ax2.set_title('TSNE Dimensions 1 & 3')

ax3 = fig.add_subplot(133)
ax3.scatter(wide_tsne[:,1], wide_tsne[:,2], color=colors, alpha=opacity)
ax3.set_title('TSNE Dimensions 2 & 3')

plt.show()

# Assuming sequence_names aligns with your wide_tsne data
mask = [seq in highlighted_sequences for seq in sequence_names]

# Filtered t-SNE data
filtered_tsne = wide_tsne[mask, :]

fig = plt.figure(figsize=(14, 4))

# Only plot the points that are in highlighted_sequences
ax1 = fig.add_subplot(131)
ax1.scatter(filtered_tsne[:, 0], filtered_tsne[:, 1], color='blue')
ax1.set_title('TSNE Dimensions 1 & 2')

ax2 = fig.add_subplot(132)
ax2.scatter(filtered_tsne[:, 0], filtered_tsne[:, 2], color='blue')
ax2.set_title('TSNE Dimensions 1 & 3')

ax3 = fig.add_subplot(133)
ax3.scatter(filtered_tsne[:, 1], filtered_tsne[:, 2], color='blue')
ax3.set_title('TSNE Dimensions 2 & 3')

plt.show()

## Make the plot
## https://shravan-kuchkula.github.io/nyc-airbnb-tsne-visual/#plotly-express-to-visualize-the-data

fig = px.scatter_3d(tsne_df, x='TSNE1', y='TSNE2', z='TSNE3',
                    hover_name=tsne_df.index, template='plotly', opacity=0.9,
                    title='Visualising sequence diversity of signal peptides (PAM distance embedded)',
                    labels={'TSNE1': 'X', 'TSNE2': 'Y', 'TSNE3':'Z'}, )

fig.write_html('scatter-3d.html')

files.download('scatter-3d.html')

fig
```
